# Structural maturation of SYCP1-mediated meiotic chromosome synapsis through conformational remodelling by molecular adapter SYCE3

**DOI:** 10.1101/2022.03.06.483192

**Authors:** James H. Crichton, James M. Dunce, Orla M. Dunne, Lucy J. Salmon, Paul S. Devenney, Jennifer Lawson, Ian R. Adams, Owen R. Davies

## Abstract

In meiosis, a supramolecular protein structure, the synaptonemal complex (SC), assembles between homologous chromosomes to facilitate their recombination. Mammalian SC formation is thought to involve hierarchical zipper-like assembly of an SYCP1 protein lattice that recruits stabilising central element (CE) proteins as it extends. Here, we combine biochemical approaches with separation-of-function mutagenesis in mice to uncover that, rather than stabilising the SYCP1 lattice, the CE protein SYCE3 actively remodels this structure during synapsis. We find that SYCP1 tetramers undergo conformational change into 2:1 heterotrimers upon SYCE3-binding, removing their assembly interfaces and disrupting the SYCP1 lattice. SYCE3 then establishes a new lattice by its self-assembly mimicking the role of the disrupted interface in tethering together SYCP1 dimers. SYCE3 also interacts with CE complexes SYCE1-SIX6OS1 and SYCE2-TEX12, providing a mechanism for their recruitment. Thus, SYCE3 remodels the SYCP1 lattice into a CE-binding integrated SYCP1-SYCE3 lattice to achieve long-range synapsis by a mature SC.

## Introduction

In meiosis, haploid germ cells are formed through the segregation of homologous chromosomes following their genetic exchange by crossing over. This requires a supramolecular protein structure, the synaptonemal complex (SC), which binds homologous chromosomes together, in synapsis, to facilitate recombination (Hunter, 2015; Zickler and Kleckner, 2015). SC assembly is directed by the inter-homologue alignments established at recombination intermediates formed at sites of induced double-strand breaks (DSBs) (Romanienko and Camerini-Otero, 2000). The mature SC structure then provides the necessary three-dimensional framework for DSB repair and crossover formation (Hunter, 2015; Zickler and Kleckner, 2015). The structural integrity of the SC is essential for meiosis across eukaryotes (Zickler and Kleckner, 2015), and SC defects are associated with human infertility, miscarriage and aneuploidy (Fan et al., 2021; Geisinger and Benavente, 2016; Schilit et al., 2020). However, the mechanism of mammalian SC assembly remains poorly understood.

The SC is a ribbon-like structure of up to 24 μm length in humans (Solari, 1980), which assembles between aligned chromosome axes at typically 400 nm initial separation, and brings their parallel axes into 100 nm synapsis (Hunter, 2015; Zickler and Kleckner, 2015). Mammalian SC assembly is thought to occur via a hierarchical zipper-like mechanism (Cahoon and Hawley, 2016; Fraune et al., 2012). Firstly, SYCP3-containing axial/lateral elements assemble along individual unaligned chromosome axes, which subsequently become aligned in homologous pairs by recombination. SYCP1 transverse filaments then assemble between aligned axes, organised with the C-termini of this coiled-coil protein within the lateral elements, and its N-termini within a midline central element (CE) (Figure 1a) (de Vries et al., 2005; Schucker et al., 2015). Head-to-head interactions between SYCP1’s N-termini are reinforced by recruitment of CE proteins SYCE3, SYCE1-SIX6OS1, and finally SYCE2-TEX12, which confer stability to the SC and allow its extension along the chromosome axis to achieve full synapsis. This hierarchical zipper-like model for SC assembly is supported by analysis of mice carrying mutations in these SC proteins, which exhibit defects at the expected stages of SC assembly with failure to recruit downstream SC proteins, and resultant chromosome asynapsis, spermatocyte death and infertility in males (Bolcun-Filas et al., 2007; Bolcun-Filas et al., 2009; de Vries et al., 2005; Gomez et al., 2016; Hamer et al., 2006; Hamer et al., 2008; Schramm et al., 2011).

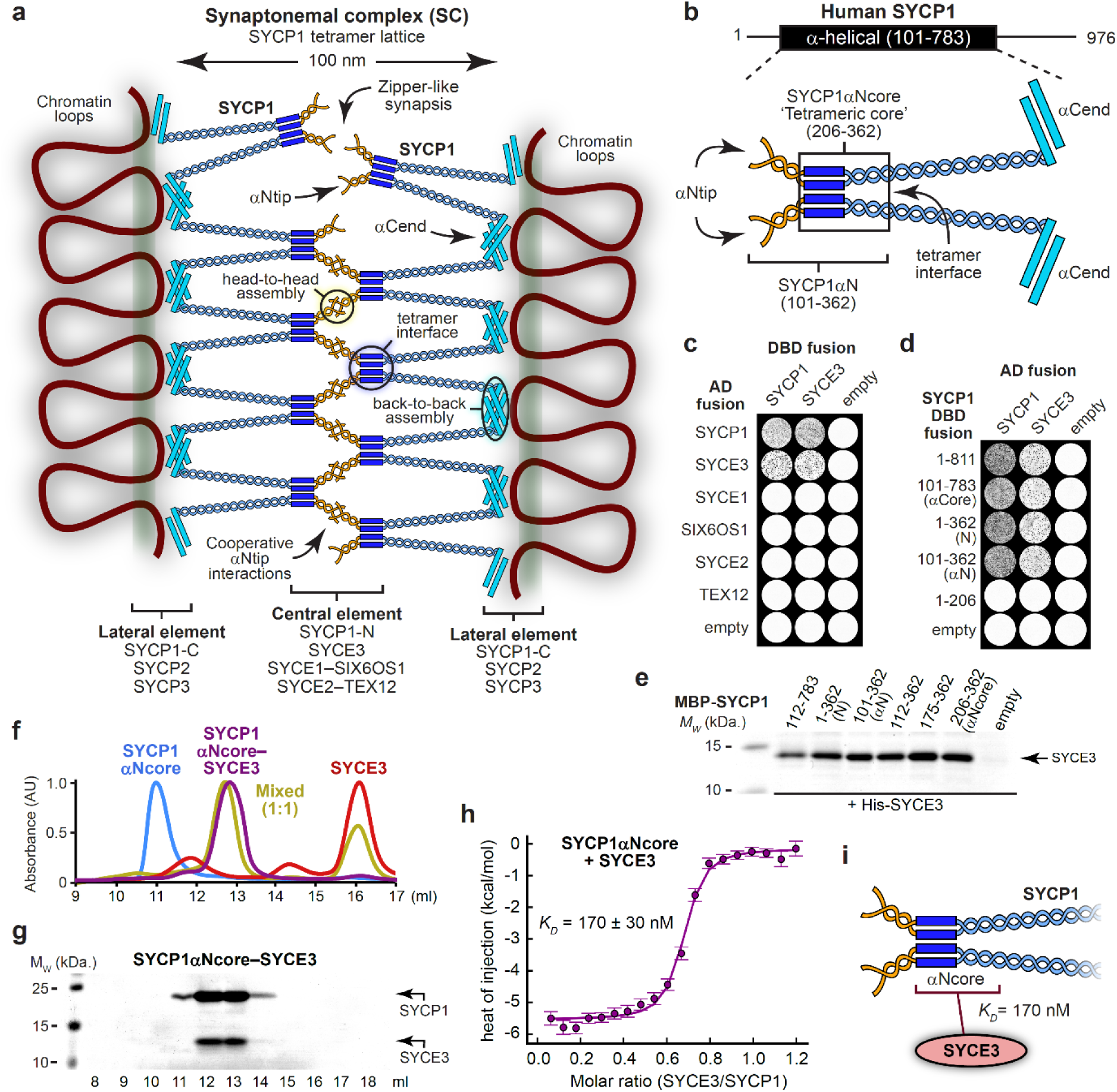
SYCP1 interacts with central element protein SYCE3. (**a**) Mammalian SC structure is defined by a supramolecular SYCP1 tetramer lattice, in which tetramer interfaces bind together parallel SYCP1 dimers and support cooperative head-to-head interactions between αNtip sites of bioriented SYCP1 tetramers, which are anchored to chromosome axes through back-to-back assembly of their α-helical C-termini (Dunce et al., 2018). (**b**) Schematic of SYCP1’s α-helical core (amino-acids 101-783), highlighting its αNtip, αCend and tetramer interface. SYCP1αNcore (amino-acids 206-362; boxed) corresponds to the tetrameric core, whereas SYCP1αN (amino-acids 101-362) is extended to include αNtips that mediate higher-order assembly. (**c,d**) Yeast two-hybrid (Y2H) analysis of (**c**) SYCP1 and SYCE3 interactions with SC proteins, and (**d**) SYCE3 interactions with truncated SYCP1 constructs. (**e**) Amylose pull-downs of SYCE3 following recombinant co-expression with MBP-SYCP1 constructs and free MBP (empty); uncropped gels are shown in Supplementary Figure 1a. (**f,g**) Size-exclusion chromatography of SYCP1αNcore (blue), SYCE3 (red), SYCP1αNcore-SYCE3 (purple) and an equimolar mixture of SYCP1αNcore and SYCE3 (yellow), shown as (**f**) UV absorbance (280 nm) chromatograms normalised to the same maximum peak height and (**g**) SDS-PAGE of SYCP1αNcore-SYCE3 elution fractions; all elution profiles are shown in Supplementary Figure 1c. (**h**) Isothermal calorimetry (ITC) of SYCE3 titrated into SYCP1αNcore, demonstrating an apparent affinity of 170 ± 30 nM (mean ± SEM, n=3). Replicates are shown in Supplementary Figure 2. (**i**) Schematic illustrating that SYCE3 binds with nanomolar affinity (*K_D_* = 170 nM) to SYCP1’s tetrameric core.

Structural and biochemical analyses have generated significant insight into the molecular mechanisms and protein interactions at play within the SC (Dunce et al., 2018; Dunce et al., 2021; Dunne and Davies, 2019a, b; Sanchez-Saez et al., 2020; Syrjanen et al., 2014). The SC’s underlying midline architecture is thought to be provided by an ‘SYCP1 tetramer lattice’ that can self-assemble *in vitro* and is stabilised by CE proteins *in vivo* (Dunce et al., 2018). In this lattice, SYCP1’s α-helical core (amino-acids 101-783) has a tetrameric structure, in which two parallel SYCP1 dimers are bound together via a ‘tetramer interface’, located towards their N-termini, within a region defined as the αNcore (amino-acids 206-362) (Figure 1b). These bifurcating molecules span between central and lateral elements, where they self-assemble through head-to-head interactions of their N-terminal α-helical tips (αNtips; amino-acids 101-111), and back-to-back interactions between DNA-binding C-termini (Figure 1a,b). Individual αNtip interactions are weak, likely to enable synaptic adjustment, meaning that head-to-head interactions depend on the cooperativity afforded through the tethering together of adjacent αNtip dimers into a lattice structure by the tetramer interface (Figure 1a,b). Thus, αNtip interactions and the tetramer interface combine into an ‘SYCP1 tetramer lattice’ that binds together chromosome axes and seemingly defines the midline structure of the SC (Figure 1a,b) (Dunce et al., 2018).

Whilst SYCP1 is sufficient for SC-like lattice assembly *in vitro* (Dunce et al., 2018; Ollinger et al., 2005), the formation of a structurally and functionally mature SC is entirely dependent on recruitment of CE proteins SYCE3, SYCE1-SIX6OS1 and SYCE2-TEX12 *in vivo* (Bolcun-Filas et al., 2007; Bolcun-Filas et al., 2009; Gomez et al., 2016; Hamer et al., 2008; Schramm et al., 2011). It has been proposed that CE proteins stabilise short- and long-range interactions within the SYCP1 lattice, including through their self-assembly (Dunce et al., 2018; Fraune et al., 2012). Indeed, SYCE3 is a dimer that self-assembles through hierarchical end-on and lateral interactions of its coiled-coil structure (Dunne and Davies, 2019a), and is thought to stabilise short-range interactions of synapsis (Schramm et al., 2011). Further, SYCE2-TEX12 self-assembles into micrometre fibres that are thought to constitute the backbone of the SC, supporting its longitudinal growth along the chromosome length (Bolcun-Filas et al., 2007; Dunce et al., 2021; Hamer et al., 2008). However, it remains unknown how CE proteins interact and integrate with the SYCP1 tetramer lattice to drive its extension along the chromosome length during SC assembly *in vivo*.

Here, we combine *in vitro* biochemical and structural studies, with genetics and imaging analysis of a separation-of-function mouse mutation *in vivo*, to uncover that SYCE3 has an essential role in actively remodelling SYCP1 tetramer lattices during the early stages of synapsis. We find that SYCE3-binding directly competes with SYCP1’s tetramer interface, disrupting the SYCP1 tetramer lattice. SYCE3 self-assembly then compensates for the disrupted interface by supporting formation of a new integrated SYCP1-SYCE3 lattice. Further, SYCE3 binds directly to CE complexes SYCE1-SIX6OS1 and SYCE2-TEX12, providing a means for their recruitment. Thus, SYCE3 acts as a molecular adapter, remodelling the nascent SYCP1 tetramer lattice into a CE-binding integrated SYCP1-SYCE3 lattice to achieve structural and functional maturation of the mammalian SC.

## Results

### The tetrameric core of SYCP1 binds to SYCE3

How do nascent SYCP1 assemblies become stabilised and extended into a mature SC? We screened for interactions between SYCP1 and individual CE proteins by yeast two-hybrid (Y2H), revealing that SYCP1 binds to SYCE3 (Figure 1c,d). This agrees with a separate study (Hernandez-Hernandez et al., 2016), and is consistent with SYCE3’s early role in hierarchical SC assembly (Schramm et al., 2011). We confirmed this interaction by pull-down of recombinant proteins and identified the SYCE3-binding site of SYCP1 as its tetrameric core (amino-acids 206-362, herein referred to as SYCP1αNcore; Figure 1e and Supplementary Figure 1a). SYCP1αNcore lies at the N-terminal end of SYCP1’s parallel coiled-coil dimers, and includes the tetramer interface that binds them together, but lacks the upstream αNtip sites that are necessary for head-to-head assembly (Figures 1b). Co-expressed SYCP1αNcore and SYCE3 purified as a stable complex that co-migrated as a distinct single species on size-exclusion chromatography (Figure 1f,g and Supplementary Figure 1b,c). Isothermal calorimetry (ITC) of the same complex, formed by mixing its purified components (Figure 1f and Supplementary Figure 1c), indicated a binding affinity of 170 ± 30 nM (Figure 1h and Supplementary Figure 2). Thus, SYCP1’s tetrameric core binds with nanomolar affinity to SYCE3, implicating this region of SYCP1 in hierarchical assembly of the SC central element (Figure 1i).

### The SYCP1 tetramer interface is disrupted by SYCE3

We expected that SYCE3 would stabilise the SYCP1 tetramer interface to support a combined role with αNtip sites in tetramer lattice formation. In contrary, size-exclusion chromatography multi-angle light scattering (SEC-MALS) identified that SYCP1αNcore-SYCE3 is a 2:1 hetero-trimer (Figure 2a and Supplementary Figure 3a). This indicates that SYCP1αNcore and SYCE3 are remodelled from tetramers and dimers upon interaction, with SYCE3-binding disrupting the SYCP1 tetramer interface. SYCP1’s full α-helical core, in which αNtips were deleted to prevent lattice assembly (amino-acids 112-783; Figure 1b), was similarly remodelled from a tetramer to a 2:1 complex by SYCE3-binding (Supplementary Figure 3b). Thus, disruption of the tetramer interface within SYCP1αNcore represents the structural consequence of SYCE3-binding to the wider SYCP1 molecule.

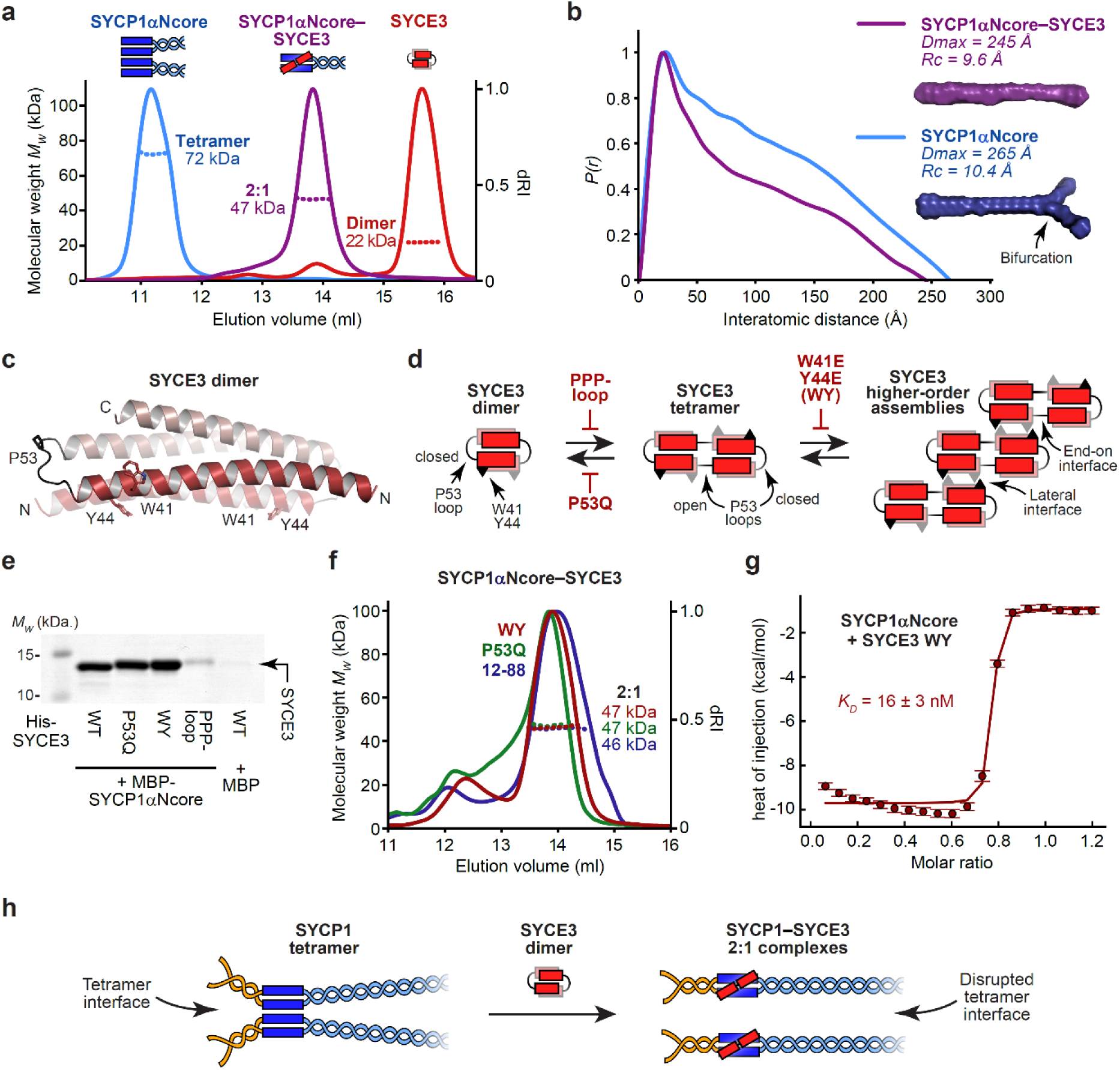
SYCP1’s tetramer interface is disrupted upon 2:1 complex formation with SYCE3. (**a**) SEC-MALS analysis showing differential refractive index (dRI; solid lines) profiles with fitted molecular weights (*Mw*; dashed lines) across elution peaks. SYCP1αNcore is a 72 kDa tetramer (blue), SYCE3 is a 22 kDa dimer (red) and SYCP1αNcore-SYCE3 is a 47 kDa 2:1 complex (purple); theoretical – 76 kDa, 21 kDa and 49 kDa. (**b**) SEC-SAXS *P(r)* distributions of SYCP1αNcore (blue) and SYCP1αNcore-SYCE3 (purple); maximum dimensions (*Dmax*) and cross-sectional radii (*Rc*) are shown alongside *ab initio* models. (**c**) Crystal structure of the SYCE3 dimer (pdb accession 6H86; Dunne and Davies (2019a); Lu et al. (2014)), in which two helix-loop-helix chains are interlaced in a four-helical bundle; amino-acids W41 and Y44, and the closed P53 loop are highlighted. (**d**) SYCE3 self-assembles into an end-on tetramer by P53 loop-opening (promoted by P53Q and inhibited by PPP-loop mutations), and into higher-order species through W41/Y44 lateral interactions (inhibited by W41E Y44E mutation; herein referred to as WY) (Dunne and Davies, 2019a). (**e**) SYCE3-binding by SYCP1 following co-expression and purification by amylose, ion exchange and size-exclusion chromatography, for MBP-SYCP1αNcore with SYCE3 point-mutations. Uncropped gel is shown in Supplementary Figure 4b. (**f**) SEC-MALS analysis showing that SYCP1αNcore forms 2:1 complexes of 47 kDa, 48 kDa and 46 kDa with SYCE3 WY, P53Q and 12-88; theoretical – 49 kDa. (**g**) ITC of SYCE3 WY titrated into SYCP1αNcore, demonstrating an apparent affinity of 16 ± 3 nM (mean ± SEM, n=3). Replicates are shown in Supplementary Figure 5a. (**h**) Schematic of SYCP1-SYCE3 2:1 complex formation. SYCP1 tetramers and SYCE3 dimers undergo conformational change, in which SYCE3 chains adopt extended open-loop conformations that bind to SYCP1 dimers, competitively inhibiting SYCP1’s tetramer interface.

How is SYCP1 remodelled into a 2:1 complex upon SYCE3-binding? The SYCP1αNcore tetramer and 2:1 complex are almost entirely α-helical and have similar melting temperatures (Supplementary Figure 3c,d), indicating their comparable structural stability, consistent with them both having biological roles. Small-angle X-ray scattering (SAXS) determined that both species have rod-like geometries, which are consistent with the theoretical 240 Å length of an extended SYCP1αNcore coiled-coil, and the known cross-sectional radii of tetrameric and trimeric coiled-coils, respectively (Figure 2b and Supplementary Figure 3e-g). Thus, SYCE3-binding remodels SYCP1αNT from an extended tetrameric coiled-coil into an extended 2:1 hetero-trimeric coiled-coil of equivalent structural stability.

SYCE3 is a dimer of helix-loop-helix chains locked in a four-helical conformation, which self-assembles into higher-order structures (Figure 2c,d) (Dunne and Davies, 2019a; Lu et al., 2014). As both helices are required for SYCP1-binding (Supplementary Figure 4a), we wondered whether SYCE3’s helix-loop-helix conformation is retained within the 2:1 complex. In SYCE3 self-assembly, chains are remodelled from helix-loop-helix to extended α-helical conformations – promoted by P53Q mutation and blocked by ‘PPP-loop’ mutation – such that they bridge between end-on dimers in a ‘domain-swap’ fashion (Figure 2d) (Dunne and Davies, 2019a). The resultant SYCE3 tetramers then assemble into higher-order structures through lateral interactions that are blocked by a W41E Y44E (WY) mutation (Figure 2d) (Dunne and Davies, 2019a). 2:1 complex formation was retained in P53Q and WY mutants, with SYCE3 WY binding to SYCP1αNcore with higher affinity than wild-type (*K_D_* = 16 ± 3 nM) (Figure 2g and Supplementary Figure 5a). In contrast, the interaction was abrogated by the PPP-loop mutation (Figure 2e,f and Supplementary Figure 5b). Thus, our mutational analysis indicates that the SYCE3 chain within the 2:1 complex adopts the extended α-helical conformation that supports SYCE3 self-assembly, rather than the helix-loop-helix conformation of the SYCE3 dimer.

In summary, SYCE3-binding and SYCP1 tetramer formation are mutually exclusive. SYCE3 forms a hetero-trimeric coiled-coil with SYCP1αNcore, competitively inhibiting the tetramer interface, and thereby remodelling an SYCP1 tetramer into SYCE3-bound dimers (Figure 2h). Hence, SYCE3 disrupts rather than stabilises SYCP1 tetramers, so is predicted to inhibit SYCP1 tetramer lattice formation.

### SYCP1-SYCE3 undergoes αNtip-mediated head-to-head assembly

SYCP1αNcore is restricted to forming a tetramer as it lacks the αNtip sites that mediate head-to-head assembly. In contrast, SYCP1’s full α-helical N-terminal region (amino-acids 101-362, herein referred to as SYCP1αN; Figure 1b) contains both αNtips and the tetramer interface, so undergoes higher-order assembly (representing tetramer lattice formation) *in vitro* (Dunce et al., 2018). Thus, to test whether SYCE3 inhibits SYCP1 tetramer lattice formation, we analysed the complex between SYCP1αN and SYCE3. SEC-MALS determined that co-expressed SYCP1αN-SYCE3 formed higher-order assemblies, in a range of molecular weights up to those observed for SYCP1αN in isolation (Figure 3a). Further, upon titration into SYCP1αN, SYCE3 was recruited to SYCP1αN assemblies, with little disruption at ten-fold molar excess (Supplementary Figure 6a). Higher-order assembly was blocked upon deletion of αNtip or introduction of mutation V105E L109E that disrupts αNtip head-to-head interactions (Dunce et al., 2018), with SYCP1αN-SYCE3 restricted to a stable 2:1 complex (Figure 3a and Supplementary Figure 6a). Thus, αNtip-mediated higher-order assembly of SYCP1αN is retained upon SYCE3-binding.

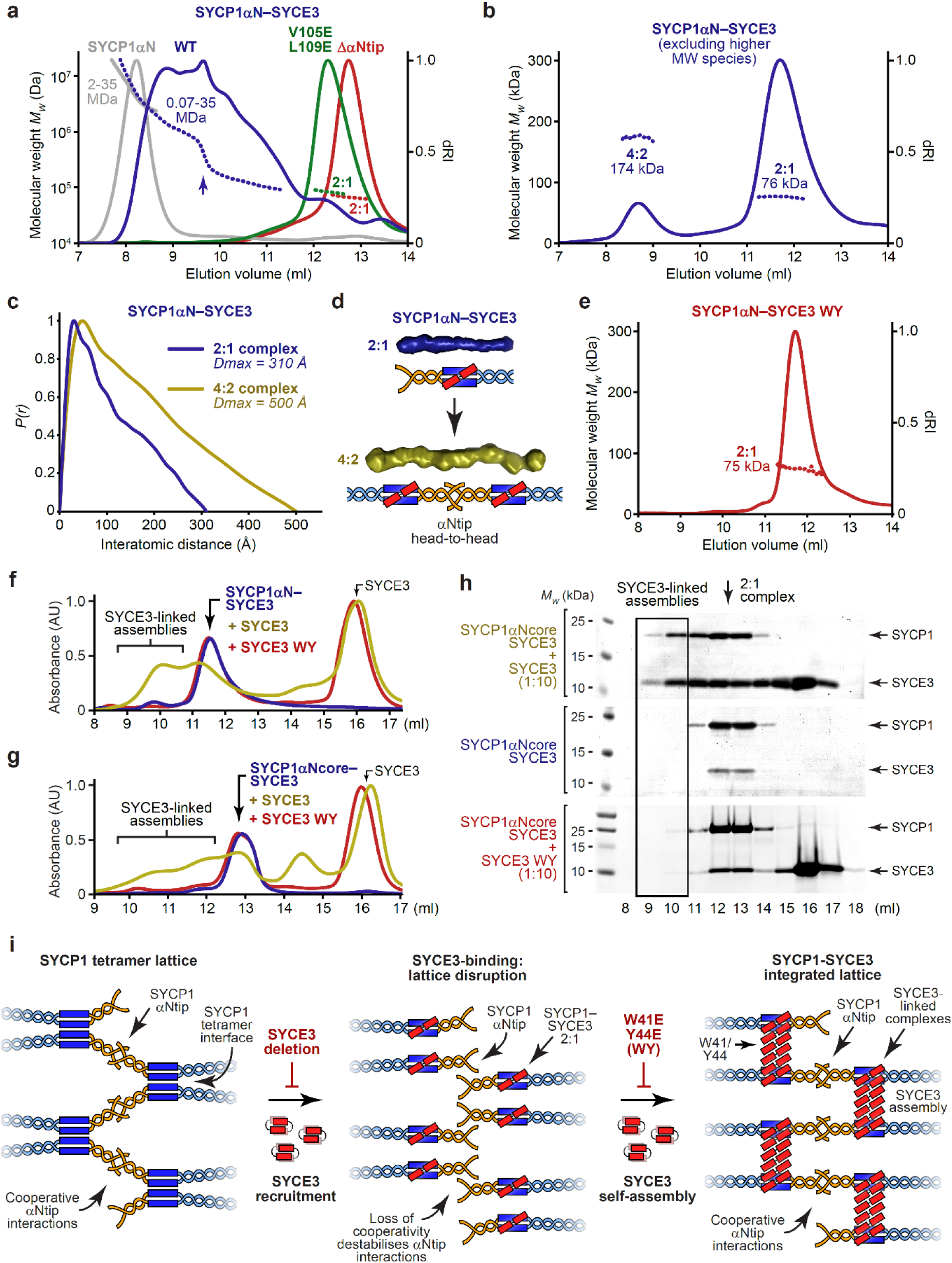
SYCP1-SYCE3 forms an integrated lattice through SYCE3 self-assembly. (**a,b**) SEC-MALS analysis. (**a**) SYCP1αN-SYCE3 (blue) forms large molecular weight species of 0.07-35 MDa (with a step from 200 kDa to 0.8 MDa indicated by an arrow), which are restricted to 2:1 complexes of 70 kDa and 75 kDa by SYCP1 ΔNtip (green) and SYCE3 WY (red) mutations, respectively; theoretical – 71 kDa and 74 kDa. SYCP1αN forms 2-35 MDa species (grey). (**b**) SYCP1αN-SYCE3 (purified to exclude large molecular weight species) demonstrates discrete 2:1 and 4:2 complexes of 76 kDa and 174 kDa, respectively; theoretical – 74 kDa and 148 kDa. (**c**) SEC-SAXS *P(r)* distributions of SYCP1αN-SYCE3 2:1 (blue) and 4:2 (yellow) species, including their *Dmax* and *Rc* values, and (**d**) *ab initio* models shown alongside a schematic of 4:2 complex formation through αNtip-mediated head-to-head interaction of 2:1 molecules. (**e**) SEC-MALS analysis showing that SYCP1αN-SYCE3 WY is a 75 kDa 2:1 complex; theoretical – 74 kDa. (**f-h**) Size-exclusion chromatography of (**f**) SYCP1αN-SYCE3 and (**g,h**) SYCP1αNT-SYCE3 upon incubation with a 10-fold stoichiometric excess of SYCE3 wild-type or WY, shown as (**f,g**) UV absorbance (280 nm) chromatograms normalised to the same maximum peak height, and (**h**) SDS-PAGE of elution fractions. Additional controls and elution fractions are shown in Supplementary Figure 8a-c. (**i**) SYCP1’s tetramer lattice depends on its tetramer and αNtip head-to-head interfaces. SYCE3 recruitment initially disrupts tetramer interfaces, forming 2:1 complexes that cannot support cooperative αNtip head-to-head interactions. Further SYCE3 molecules assemble, mediated by W41 and Y44 amino-acids (inhibited by the WY mutation), into structures that incorporate and link together SYCP1-SYCE3 complexes, mimicking tetramer associations to form an integrated SYCP1-SYCE3 lattice.

If SYCP1αN-SYCE3 assembles through the same αNtip head-to-head interactions that are responsible for SYCP1 tetramer lattice assembly, then we would predict the presence of an assembly intermediate in which 2:1 species interact ‘head-to-head’ in 4:2 complexes. Accordingly, upon selective purification to remove higher-order species, we determined that the lowest molecular weight species of SYCP1αN-SYCE3 correspond to 2:1 and 4:2 complexes (Figure 3b). In support of this, the MBP-fusion complex formed only 2:1 and 4:2 species, likely owing to inhibition of higher-order assembly by steric hindrance (Supplementary Figure 6b). SAXS analysis of the 2:1 and 4:2 complexes revealed rod-like molecules in which 4:2 complexes are almost twice as long as 2:1 complexes, consistent with their formation through end-on interactions of 2:1 complexes (Figure 3c,d and Supplementary Figure 6c,d). Thus, we conclude that higher-order assembly, mediated by αNtip head-to-head interactions, is retained upon SYCE3 binding to SYCP1αN, despite disruption of the tetramer interface.

### An integrated SYCP1-SYCE3 lattice is established by SYCE3 self-assembly

How does SYCP1αN-SYCE3 undergo αNtip-mediated assembly in absence of the tetramer interface? The SYCE3 WY mutation, which blocks lateral interactions of the SYCE3 self-assembly pathway (Figure 2d), also blocked higher-order assembly and restricted SYCP1αN-SYCE3 to a 2:1 complex, despite the presence of αNtips (Figure 3e). The ability to block SYCP1αN assembly by deleting the αNtip (Dunce et al., 2018), or disrupting the tetramer interface by SYCE3 WY-binding (Figure 3e), is consistent with the tetramer interface providing the cooperativity necessary to support individually weak αNtip interactions. Further, these data suggest that SYCE3’s lateral assembly interactions must compensate for the missing tetramer interface between SYCP1 dimers by providing an analogous ‘tetramer-like’ interface between adjacent SYCE3-bound SYCP1 dimers.

The addition of free SYCE3 to SYCP1αN-SYCE3 2:1 and 4:2 complexes triggered their assembly into higher-order species in a manner that was blocked by the WY mutation (Figure 3f and Supplementary Figure 7a,b). Thus, ‘tetramer-like’ interfaces provided by SYCE3’s lateral assembly interactions likely involve SYCP1-SYCE3 2:1 complexes being linked together by additional SYCE3 molecules, rather than by direct interactions. Given the shared SYCE3 extended chain conformation, we wondered whether SYCP1-SYCE3 2:1 complexes may structurally mimic SYCE3 tetramers, allowing their incorporation as laterally-interacting units within SYCE3 assemblies. This explains disruption by the WY mutation, and predicts that ‘tetramer-like’ interfaces are independent of αNtips. Accordingly, SYCP1αNcore-SYCE3, which lacks αNtips and forms only 2:1 complexes in isolation, underwent higher-order assembly upon addition of free SYCE3, but not of the WY mutant (Figure 3g,h and Supplementary Figure 7c). Thus, SYCP1-SYCE3 forms an integrated lattice, in a similar manner to the SYCP1 tetramer lattice, through cooperativity between αNtip head-to-head interactions and ‘tetramer-like’ interfaces between SYCE3-bound SYCP1 dimers mediated by their lateral incorporation into SYCE3 assemblies (Figure 3i).

In summary, our findings suggest that SYCE3 remodels the SYCP1 tetramer lattice into a structurally distinct integrated SYCP1-SYCE3 lattice (Figure 3i). Firstly, SYCE3-binding disrupts SYCP1’s tetramer interface, dissolving the tetramer lattice into SYCP1-SYCE3 2:1 complexes. Then, SYCE3 assemblies link together 2:1 molecules, mimicking the disrupted tetramer interface, to support cooperative αNtip head-to-head interactions within an integrated SYCP1-SYCE3 lattice (Figure 3i).

### Syce3^WY/WY^ mice are infertile with failure of SC assembly

We next investigated how the ability of SYCE3 to remodel SYCP1 tetramer lattices *in vitro* relates to SC assembly *in vivo*. The SYCE3 WY mutation separates the disruptive and integrative functions of SYCE3, triggering SYCP1 tetramer lattice disruption, whilst failing to form an integrated SYCP1-SYCE3 lattice (Figure 3i). Hence, the WY mutation is predicted to be more deleterious than a simple SYCE3 deletion, in which SYCP1 tetramer lattices can be retained but cannot be remodelled (Figure 3i). Thus, to test our model for SYCP1 lattice remodelling by SYCE3, we generated and analysed SC assembly in *Syce3^WY/WY^* mice.

As *Syce3^WY^* has the potential to act in a dominant-negative manner, we circumvented the need for fertile *Syce3^WY/+^* heterozygotes by analysing *Syce3^WY/WY^* homozygotes born directly from CRISPR/Cas9 gene editing in zygotes (Figure 4a) (Teboul et al., 2017; Wang et al., 2013). We also generated control *Syce3^PAM/PAM^* mice possessing the silent protospacer adjacent motif (PAM) mutations introduced alongside the WY mutation in *Syce3^WY/WY^* mice, and *Syce3^Δ/Δ^* mice carrying *Syce3* frameshift deletions preceding or encompassing the WY mutation site as potential null alleles (Figures 4a,b and Supplementary Figure 8a,b). The *Syce3^PAM/PAM^* control mice, which encode wild-type SYCE3 protein, control for off-target CRISPR/Cas9 editing events and unexpected effects of the synonymous and non-coding PAM site mutations on *SYCE3* expression.

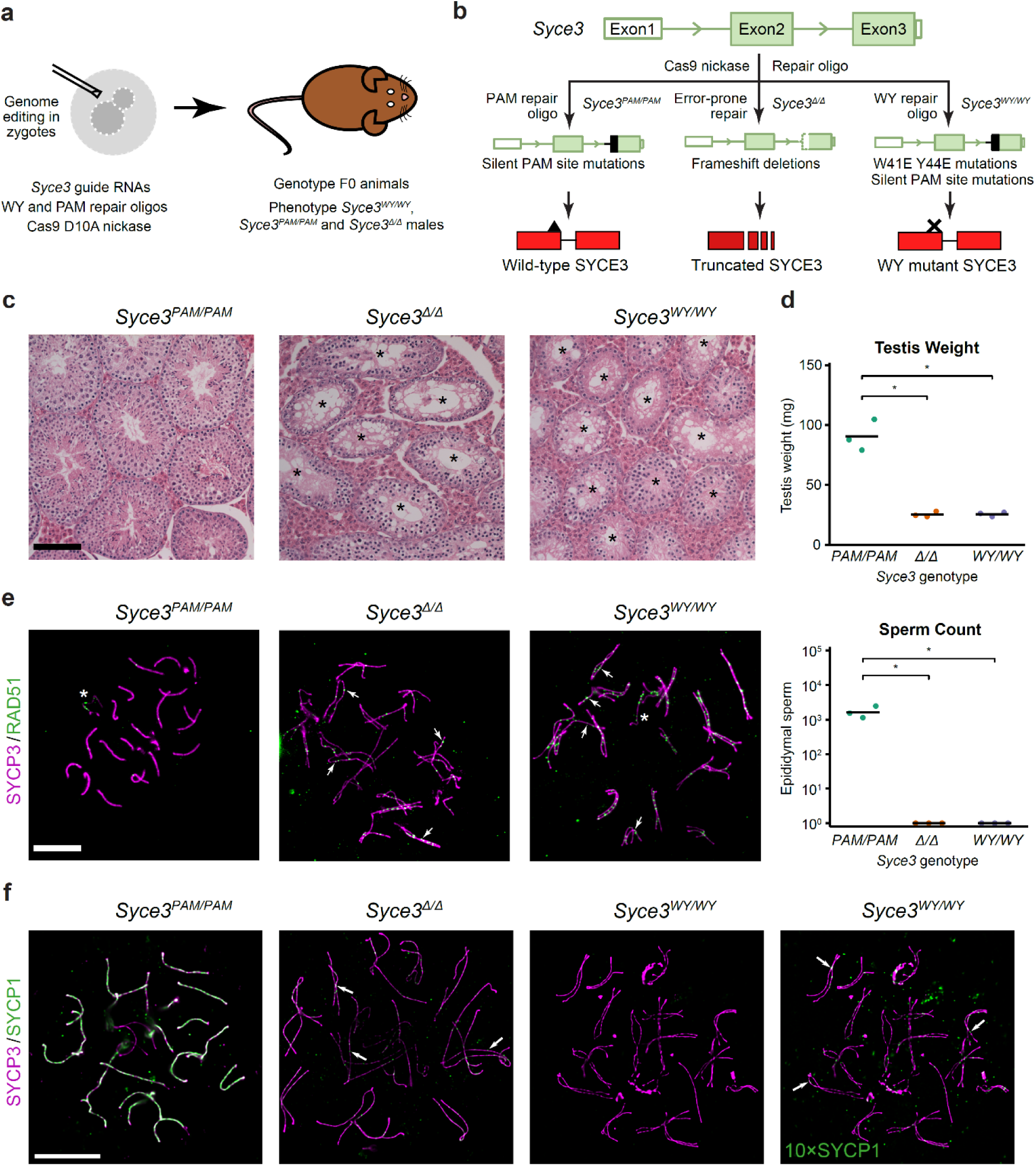
SYCE3 self-assembly is required for SC assembly and meiotic progression *in vivo*. (**a**) Generation and analysis of *Syce3* mutant mice. Zygotes were microinjected with *Syce3* CRISPR/Cas9 genome editing reagents and male F0 animals with desired genotypes analysed. (**b**) *Syce3* genome editing strategy. *Syce3* protein coding regions are filled green, regions repaired by the repair oligos are filled black, and their effects on SYCE3 protein (red) are indicated. (**c**) Haematoxylin and eosin staining of *Syce3* testes sections. Asterisks indicate tubules with a spermatogenic block and depletion of post-meiotic spermatids. Scale bar, 100 µm. (**c**) Testis weights (top) and sperm counts (bottom) of *Syce3* animals. Testis weights for each animal are shown, with the mean for each genotype indicated with a black horizontal bar. Means are 90.6, 25.4 and 25.7 mg. *, p<0.01, (Student’s t-test, n=3). Total numbers of sperm isolated from one epididymis per animal are plotted with the mean for each genotype indicated with a black horizontal bar. Means are 1709, 0 and 0 sperm. *, p<0.05 (Student’s t-test, n=3). Summary statistics are in Supplementary Table 2. (**e,f**) Widefield imaging of pachytene *Syce3^PAM/PAM^* and asynapsed pachytene *Syce3^Δ/Δ^* and *Syce3^WY/W^* meiotic chromosome spreads immunostained for SYCP3 (magenta) and either RAD51 (**e**, green) or SYCP1 (**f**, green). Examples of paired asynapsed chromosomes are indicated with arrowheads (**e**), axial SYCP1 foci with arrows (**f**) and sex chromosomes with an asterisk. Scale bar, 10 µm.

*Syce3^WY/WY^* and *Syce3^Δ/Δ^* homozygotes exhibited severely reduced testis weights and epididymal sperm counts, with testis histology indicating a severe block in meiosis, whereas *Syce3^PAM/PAM^* homozygotes had no overt defects (Figure 4c,d) (Crichton et al., 2017). Furthermore, no overt phenotypic mosaicism was detected in these animals (Figure 4c,d). Immunostained chromosome spreads demonstrated that RAD51 recombination foci formed in *Syce3^WY/WY^* and *Syce3^Δ/Δ^* spermatocytes (Figure 4e), but extensive regions of SYCP3-coated chromosome axes did not synapse and SYCP1 recruitment was fragmented (Figure 4f), with meiosis arrested at pachytene (Supplementary Figure 8c). These findings correlate with the reported *Syce3^-/-^* phenotype (Schramm et al., 2011), confirming that SYCE3’s W41/Y44-mediated lateral self-assembly interactions are essential for *Syce3* function and SC assembly *in vivo*. Thus, the nature and severity of the *Syce3^WY/WY^* phenotype are consistent with our biochemical findings and support our model for SC assembly through SYCP1 lattice remodelling by SYCE3.

### SYCP1 assembly is more severely disrupted in Syce3^WY/WY^ than Syce3^Δ/Δ^ mice

Although fragmented SYCP1 staining was detected on chromosome axes in both *Syce3* mutants, SYCP1 staining was considerably less prominent in *Syce3^WY/WY^* than *Syce3^Δ/Δ^* nuclei, requiring a 10-fold increase in image brightness for detection (Figure 4f). This more deleterious effect of *Syce3^WY^* than *Syce3* null alleles on SYCP1 assembly presumably reflects the ability of SYCE3 to disrupt the SYCP1 tetramer lattice. We therefore examined their SYCP1 foci in detail using structured illumination microscopy (SIM). In *Syce3^PAM/PAM^* pachytene nuclei, SYCP1 localised between pairs of SYCP3-stained axes, often appearing as chains of doublet foci (consistent with its biorientation), with occasional discontinuities, and sometimes as linear extensions tightly associated with one of the axes (Figure 5a and Supplementary Figure 9a). In asynapsed *Syce3^WY/WY^* and *Syce3^Δ/Δ^* pachytene nuclei, SYCP1 staining was less orderly, consisting of mostly axial foci (within 35 nm of the SYCP3 axis; Supplementary Figure 9b), and we rarely observed doublet foci linking paired axes (Figure 5a-c). In agreement with widefield imaging, the number and intensity of both axial and non-axial SYCP1 foci were far greater in *Syce3^Δ/Δ^* than *Syce3^WY/WY^* nuclei (Figure 5c,d and Supplementary Figure 9b-e). Thus, the SYCP1 foci present in *Syce3^Δ/Δ^* nuclei likely include SYCP1 tetramer lattices whose assembly and/or stability are actively disrupted by the SYCE3 WY mutation.

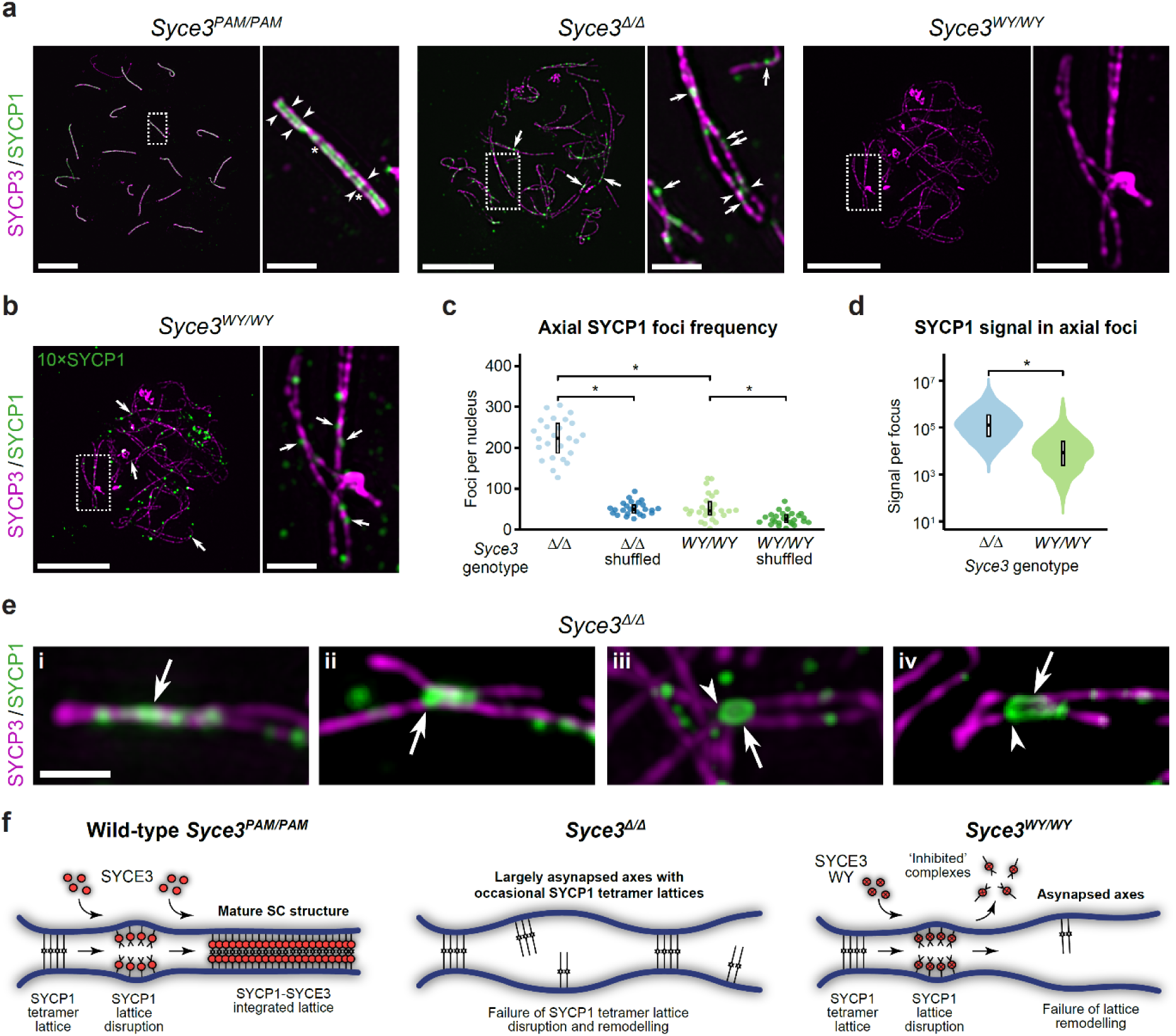
SYCP1 tetramer lattices are disrupted by SYCE3 WY *in vivo*. (**a,b**) SIM images of pachytene *Syce3^PAM/PAM^* and asynapsed pachytene *Syce3^Δ/Δ^* and *Syce3^WY/WY^* meiotic chromosome spreads immunostained for SYCP3 (magenta) and SYCP1 (green). The brightness of the SYCP1 channel in the *Syce3^WY/W^* image in (a) has been increased ten-fold to generate the image in (b). Example axial SYCP1 foci (arrows), SYCP1 foci doublets (arrowheads), and SYCP1 discontinuities (asterisks) are indicated. Example linear SYCP1 extensions are shown in Supplementary Figure 10a. Scale bars, 10 µm for low magnification images, 1 µm for enlarged regions. (**c**) SYCP1 focus centroids within 35 nm of the SYCP3 axis mask in *Syce3* asynapsed pachytene nuclei were classed as axial (Supplementary Figure 9b) and counted. Axial foci in shuffled datasets represent the 95% upper bound after twenty rounds of randomly assigning each SYCP1 focus centroid a location within that nucleus. Crossbars represent quartiles; *, p<0.01 (Mann-Whitney U test, medians are 223, 49, 46 and 25 foci); 3 animals analysed for each *Syce3* genotype. Summary statistics are in Supplementary Table 2 (**d**) The total SYCP1 signal in each axial SYCP1 focus in (a) was measured. Crossbars represent quartiles; *, p<0.01 (Mann-Whitney U test, medians are 123441 and 8485 arbitrary units); 3 animals analysed for each *Syce3* genotype. Data for individual animals are shown in Supplementary Figure 9c. Summary statistics are in Supplementary Table 2 (**g**) Summary of the consequence of *Syce3* mutations. In wild-type and *Syce3^PAM/PAM^*, nascent SYCP1 tetramer lattices are remodelled by wild-type SYCE3 protein into integrated SYCP1-SYCE3 lattices of mature SC structure. In *Syce3^Δ/Δ^*, nascent SYCP1 tetramer lattices are retained by not matured owing to the absence of SYCE3. In *Syce3^WY/WY^*, nascent SYCP1 tetramer lattices are disrupted but cannot be remodelled by SYCE3 WY, leaving largely undecorated axes.

We next investigated whether the SYCP1 tetramer lattices in *Syce3^Δ/Δ^* spermatocytes resemble mature SCs, in which chains of doublet SYCP1 foci bridge between synapsed axes (Figure 5a), by focussing on large extended SYCP1 assemblies at sites of close proximity between paired SYCP3 axes. In some cases, these large extended SYCP1 foci consisted of linear structures associated with tightly apposed SYCP3 axes, or were assembled in the gap between pairs of SYCP3 axes, but did not appear to consist of doublet foci (Figure 5e, panel i-ii). In other cases, the large extended SYCP1 foci comprised of pairs of linear SYCP1 assemblies, connected to each other by terminal doublet-like structures, but extending separately along each axis (Figure 5e, panels iii-iv). Whilst these terminal doublet-like structures appear to be unable to develop into chains in the absence of SYCE3, their extending linear SYCP1 assemblies resemble the linear extensions of SYCP1 observed in some regions of the mature SC (Supplementary Figure 9a). Thus, SYCP1 tetramer lattices can contribute to the doublet-like foci between closely paired SYCP3 axes, but lattice remodelling by SYCE3 is required for their extension into the doublet chains of the mature SC.

In summary, *Syce3* mutants exhibit defects at distinct SC assembly stages that support our biochemical findings and model for SYCP1 lattice remodelling by SYCE3. Firstly, *Syce3^Δ/Δ^* captures the formation of SYCP1 tetramer lattices between axes, which cannot be remodelled in absence of SYCE3, so fail to develop into a mature SC (Figure 5f). Secondly, *Syce3^WY/WY^* captures the stage at which SYCP1 tetramer lattices are disrupted by SYCE3 but cannot be remodelled into integrated SYCP1-SYCE3 lattices (Figure 5f). Finally, in wild-type and control *Syce3^PAM/PAM^* mice expressing wild-type SYCE3 protein, SYCE3 remodels SYCP1 tetramer lattices into integrated SYCP1-SYCE3 lattices that support full SC maturation (Figure 5f).

### SYCE3 recruits SYCE1-SIX6OS1 and SYCE2-TEX12 complexes

What is the functional consequence of SYCE3 integration into the SYCP1 lattice? The CE contains three high-affinity ‘building-block’ complexes: SYCP1-SYCE3 (this study), SYCE1-SIX6OS1 (Sanchez-Saez et al., 2020) and SYCE2-TEX12 (Dunce et al., 2021). We identified through biochemical pull-downs that SYCE3 binds to SYCE1-SIX6OS1 and SYCE2-TEX12 complexes (Figure 6a-d and Supplementary Figures 10a-c and 11a). The SYCE1-SIX6OS1 interaction is mediated by SYCE3’s N-terminus binding to SYCE1’s α-helical core (Figure 6a,b and Supplementary Figure 10a,b), whilst the SYCE2-TEX12 interaction requires the presence of both SYCE2 and TEX12 components (Figure 6d and Supplementary Figure 11a). In both cases, complexes largely dissociated during size-exclusion chromatography (Figure 6e and Supplementary Figures 10d and 11b), consistent with micromolar binding affinities. Indeed, microscale thermophoresis identified a binding affinity between SYCE3 and SYCE2-TEX12 of 22 ± 2 μM (Figure 6f and Supplementary Figure 11c-i). These SYCE3 interactions are two orders of magnitude weaker than constituent interactions of the SC’s core complexes, suggesting that they are dynamic, with rapid turnover of binding partners, and likely achieve stability through cooperativity within the SC lattice. Further, SYCE3 promoted fibrous assembly of SYCE2-TEX12 (Figure 6g and Supplementary Figure 12), suggesting that it functions both in the recruitment and assembly of the CE.

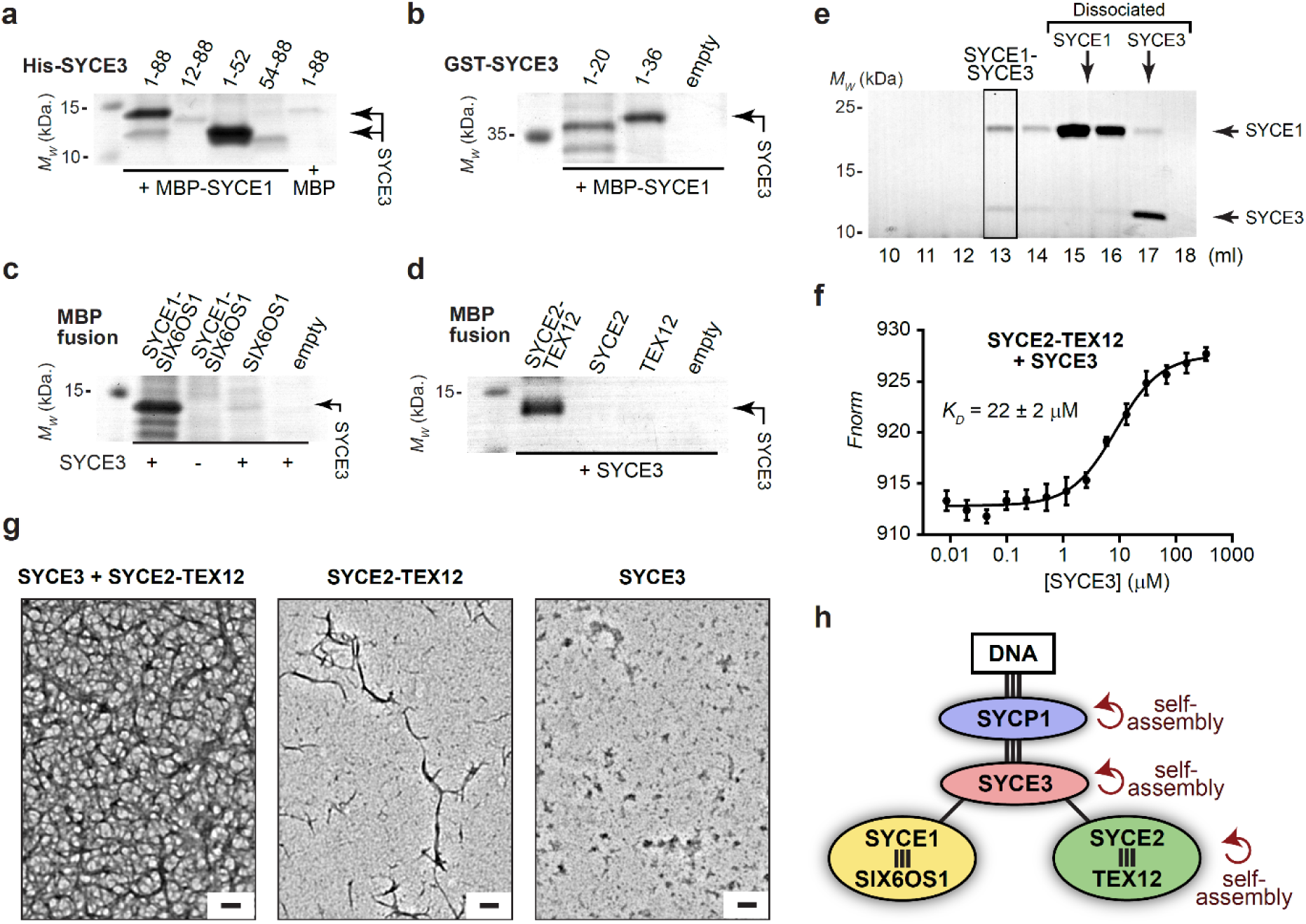
SYCE3 interacts with SYCE1-SIX6OS1 and SYCE2-TEX12 central element complexes. (**a-d**) Amylose pull-downs of (**a,c,d**) His-SYCE3 and (**b**) GST-SYCE3 and free GST (empty), following recombinant co-expression with (**a**) MBP-SYCE1core and free MBP (empty), (**b**) MBP-SYCE1core, (**c**) MBP-SIX6OS1N in the presence or absence of SYCE1core, and free MBP (empty), and (**d**) SYCE2-TEX12 core, SYCE2 core, TEX12 core and free MBP (empty). Uncropped gels are shown in Supplementary Figures 11Aa and 12a. (**e**) Size-exclusion chromatography of the SYCE1-SYCE3 complex, showing some complex retention (boxed) but mostly dissociation of components; UV chromatograms are shown in Supplementary Figure 11b. (**f**) MST of SYCE3 titrated into 150 nM SYCE2-TEX12core, demonstrating an apparent binding affinity of 21.8 ± 2.1 μM (mean ± SEM, n=3); full data are shown in Supplementary Figure 12c-i. (**g**) Electron microscopy of SYCE2-TEX12 (full-length) following incubation with a two-fold molar excess of SYCE3; individual components are shown for comparison. Scale bars, 100 nm. Full panels are shown in Supplementary Figure 13. (**h**) Interaction network of SC central element proteins, indicating strong interactions (three lines), weak interactions (single lines) and self-assembly.

Our data suggest that SYCE3 acts as a molecular adapter that binds together the CE’s ‘building-block’ complexes, through a combination of high- and low-affinity binding interfaces, and self-assembly interactions, to assemble a mature SC structure (Figure 6h). SYCE3 remodels and integrates into the SYCP1 lattice, establishing binding sites that cooperativity recruit SYCE1-SIX6OS1 and SYCE2-TEX12, and stimulate SYCP1-SYCE3 and SYCE2-TEX12 assembly, to structurally reinforce and drive SC growth (Figure 7). This model explains the failed extension of SYCP1 assemblies in *Syce3^Δ/Δ^* nuclei, and the severe disruption of SYCP1 assemblies in *Syce3^WY/WY^*. Hence, we uncover an essential role for SYCE3 in integrating the CE’s distinct architectural units into a structurally and functionally mature SC.

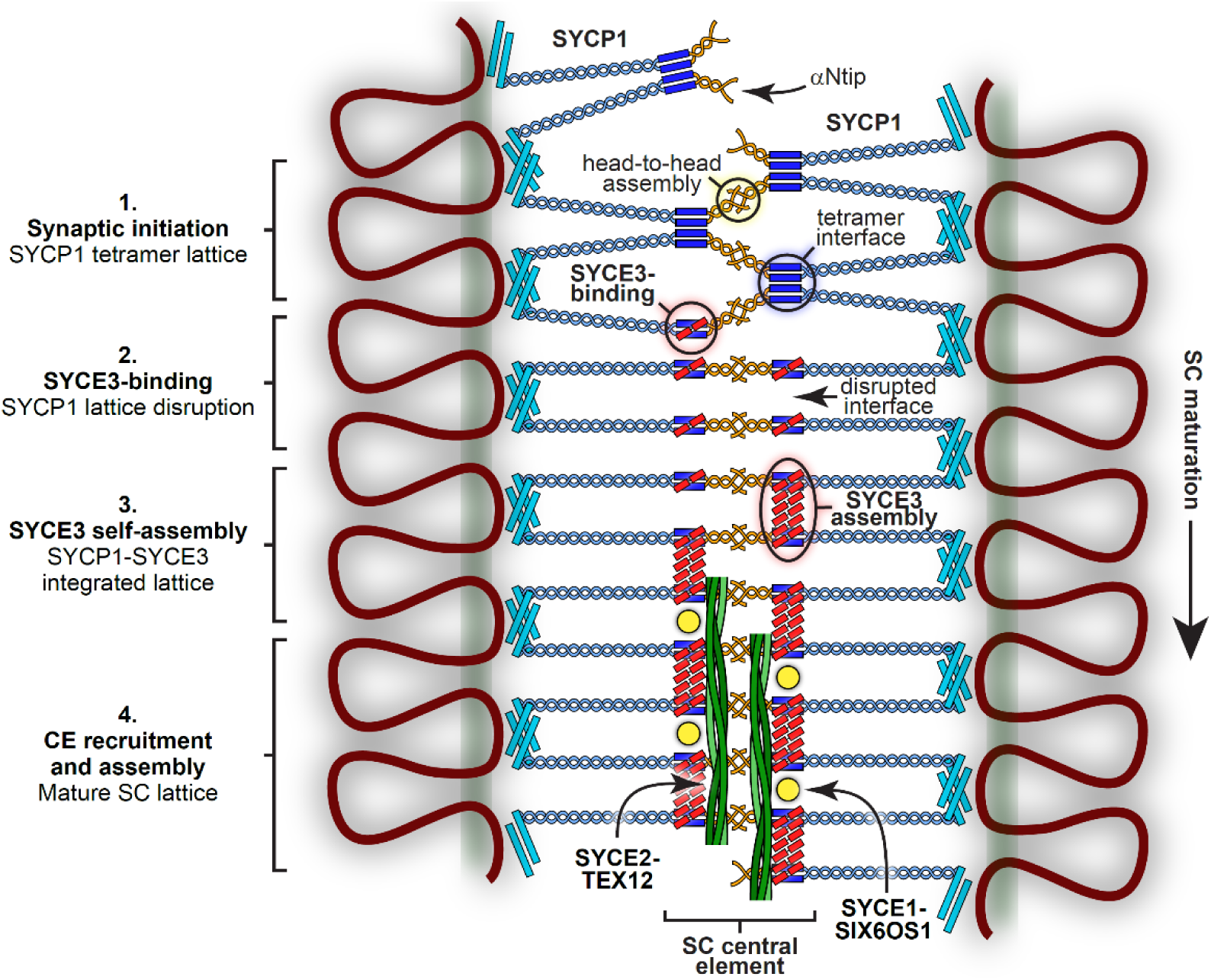
Model for SC maturation through SYCP1 lattice remodelling and integration by SYCE3. Model for the structural maturation of the SC through SYCE3-mediated remodelling of the SYCP1 lattice and recruitment of CE complexes. 1. Synaptic initiation and local lattice extension occur through the recruitment and assembly of SYCP1 tetramer lattices between chromosome axes. 2. SYCE3 recruitment disrupts the tetramer lattice by binding to SYCP1 dimers and competitively inhibiting the tetramer interface. 3. SYCE3 self-assembly then binds together SYCP1-SYCE3 complexes, mimicking the role of the tetramer interface, resulting in the remodelling of the initial SYCP1 tetramer lattice into an SYCP1-SYCE3 integrated lattice. 4. Incorporated SYCE3 assemblies recruit and initiate assembly of SYCE1-SIX6OS1 and SYCE2-TEX12 complexes that provide short-range and long-range fibrous supports that stabilise the SC’s extension along the chromosome length.

## Discussion

Our combined biochemical and separation-of-function mutagenesis studies provide a new paradigm for the role of CE protein SYCE3 in mammalian SC assembly. Firstly, rather than simply stabilising existing SYCP1 assemblies, SYCE3 remodels the SYCP1 tetramer lattice into an integrated SYCP1-SYCE3 lattice that enables SC growth (Figure 7). Secondly, SYCE3 promotes SYCP1-SYCE3 lattice extension and SYCE2-TEX12 fibre formation through SYCP1, SYCE3 and SYCE2-TEX12 self-assembly. Finally, SYCE3 is central within a network of low-affinity interactions that bind together the SC’s high-affinity heteromeric complexes SYCP1-SYCE3, SYCE1-SIX6OS1 and SYCE2-TEX12 (Figure 6h). Thus, SYCE3 performs multiple distinct roles as a molecular adapter of SC assembly.

The remodelling of SYCP1 tetramer lattices into integrated SYCP1-SYCE3 lattices involves multiple conformational remodelling and self-assembly mechanisms. Upon binding, SYCP1 and SYCE3 undergo conformational change from tetramers and dimers to a 2:1 hetero-trimeric complex, in a process that competes with, and thereby disrupts, SYCP1’s tetramer interface. In parallel, SYCE3 self-assembles by conformational domain-swap of dimers into tetramers that interact laterally (Figure 2d) (Dunne and Davies, 2019a). These SYCE3 assemblies bind to, and link together, SYCP1-SYCE3 complexes, mimicking the role of the disrupted tetramer interface to establish a new integrated SYCP1-SYCE3 lattice (Figure 7). Thus, SYCP1 and SYCE3 exhibit conformational plasticity, with the same protein sequences forming multiple distinct conformations and assemblies. The formation of alternative conformations has been observed in other coiled-coil systems, and is attributed to their similar interfaces giving rise to only small differences in the free-energy of folding (Croasdale et al., 2011; Lizatovic et al., 2016; Roder and Wales, 2017). Hence, this plasticity is likely consequential of the coiled-coil nature of SC proteins.

Are the SYCP1 tetramer lattice and integrated SYCP1-SYCE3 lattice temporally exclusive or could they co-exist within an assembled SC? The similar melting temperatures of SYCP1αNcore and its complex with SYCE3 suggest that both lattices have similar stability. Thus, the SC could be sustained locally by either lattice type, which could be dynamically and reversibly remodelled through active or reactive mechanisms such as local SYCE3 availability and post-translational modifications (Jordan et al., 2012). A more adaptive SYCP1 tetramer lattice may be required to enable synaptic adjustment of initially poorly aligned axes prior to formation of a mature SC structure, whereas the SYCP1-SYCE3 lattice may represent a more rigid structure (Figure 7). The structural heterogeneity that would result from the co-existence of both lattice types is consistent with the irregularities in SYCP1 structures observed within the assembled mouse SC by immunofluorescence (Figure 5a and Supplementary Figure 9a) and EM (Spindler et al., 2019). Further, the distinct SYCP1 tetramer and integrated SYCP1-SYCE3 lattice structures may have functional consequence, such as permitting differential access to recombination sites as a means of locally regulating meiotic recombination. This may explain the observed structural alterations of the SC at recombination sites in *C. elegans* (Libuda et al., 2013; Woglar and Villeneuve, 2018). Thus, it be of great interest to determine whether structural heterogeneity and/or dynamic structural remodelling of the SC influence recombination frequencies and crossover outcomes in mammals.

The large axial SYCP1 foci formed in *Syce3^Δ/Δ^* but not *Syce3^WY/WY^* spermatocytes likely represent SYCP1 tetramer lattices that were trapped owing to the lack of SYCE3 (Figure 5f). Whilst some assembled between paired axes at sites of potential synapsis, others were located on individual axes, raising the question of how SYCP1 tetramer lattices can assemble on single rather than paired axes. As SYCP1 assembles into tetramer lattices in absence of DNA *in vitro* (Dunce et al., 2018; Ollinger et al., 2005), one side of the lattice may bind to the axis, leaving the other free to subsequently capture the paired axis (Figure 5f). Alternatively, SYCP1 tetramer lattices may assemble between chromatin loops or sister chromatids of the same axis. Hence, SYCE3-binding may have an additional role in redirecting SYCP1 lattices towards inter-homologous synapsis. As SYCP1 assemblies on individual axes are also present in *Syce1^-/-^*, but are rare in *Syce2^-/-^* spermatocytes (Bolcun-Filas et al., 2007; Bolcun-Filas et al., 2009; Schramm et al., 2011), this likely involves the stabilising interactions of SYCE2-TEX12 proteins affording a cooperativity that strongly favours the formation of a single continuous inter-axial lattice rather than short discontinuous patches within individual axes.

The interaction network of the SC, which we defined through biochemical and biophysical analysis of recombinant SC proteins (Figure 6h) (Dunce et al., 2018; Sanchez-Saez et al., 2020), agrees with previous knockout phenotypes, co-immunoprecipitation and heterologous co-localisation studies (Bolcun-Filas et al., 2007; Bolcun-Filas et al., 2009; de Vries et al., 2005; Gomez et al., 2016; Hamer et al., 2006; Hamer et al., 2008; Hernandez-Hernandez et al., 2016; Lu et al., 2014; Schramm et al., 2011). Our findings reveal an apparent dichotomy of high-affinity (nanomolar) CE complexes – SYCP1-SYCE3, SYCE1-SIX6OS1 and SYCE2-TEX12 – that are held together by low-affinity (micromolar) interactions. This divides the SC’s interactions into long-lasting complexes that likely represent its ‘building-block’ structures, and those that are transient and rapidly exchanged within a dynamic SC assembly. Further, SYCP1, SYCE3 and SYCE2-TEX12 undergo self-assembly through the cooperative action of similarly low-affinity individual interfaces (Figure 6h) (Dunce et al., 2018; Dunce et al., 2021). Hence, SYCP1-SYCE3, SYCE1-SIX6OS1 and SYCE2-TEX12 represent the SC’s discrete heteromeric units that interact and self-assemble through low-affinity interfaces that are likely stabilised by cooperativity within the SC lattice.

SC assembly involves two distinct SYCP1 lattices, which are interconverted by SYCE3 modelling, and the binding together of its building-block complexes by low-affinity binary and self-assembly interactions. Together, these provide means for formation of a dynamic, adaptive and structurally heterogeneous SC from a series of well-defined and specific protein-protein interfaces. Indeed, the SC may be considered as having emergent functions (Pancsa et al., 2019), which could not be predicted from its individual protein components *a priori*, but are inherently defined by its constituent interactions. In this respect, active or passive remodelling of the SC may allow rapid bending, twisting and distortion of the central element in adaptation to mechanical stresses. This may influence accessibility of recombination factors to recombining DNA, and dynamically regulate the frequency, distribution and outcomes of meiotic recombination. This functionality would not be possible if the SC had an homogenous and rigid structure. Hence, the complexity of interactions that underly the SC’s structure are likely critically important to its function. Hence, the SC is one of the most intriguing and enigmatic biological structures, of which structural and functional understanding are critical to uncovering the molecular basis of meiotic recombination.

## Materials and Methods

### Recombinant protein expression and purification

SYCP1, SYCE3, SYCE1, SYCE1-SIX6OS1 and SYCE2-TEX12 protein constructs and complexes were purified as previously described (Davies et al., 2012; Dunce et al., 2018; Dunce et al., 2021; Dunne and Davies, 2019a, b; Sanchez-Saez et al., 2020). In general, proteins were expressed as His- or His-MBP fusions in BL21(DE3) *E. coli* cells (Novagen®), and purified from lysate through Ni-NTA (Qiagen) or amylose (NEB) affinity, with removal of the tag by TEV protease treatment, followed by anion exchange chromatography (HiTrap Q HP, Cytiva) and size exclusion chromatography (HiLoad^TM^ 16/600 Superdex^TM^ 200, Cytiva) in 20 mM Tris pH 8.0, 150 mM KCl, 2 mM DTT. Purified proteins were concentrated using Amicon Ultra® 10,000 MWCO centrifugal filter units (Millipore) or Microsep™ Advance 3kDa (PALL) centrifugal filter units, and flash-frozen in liquid nitrogen for storage at -80°C. Samples were analysed by Coomassie-stained SDS-PAGE, and concentrations were determined using a Cary 60 UV spectrophotometer (Agilent) with molecular weights and extinction coefficients calculated by ExPASY ProtParam (http://web.expasy.org/protparam/).

### Co-expression amylose pull-down assays

Protein-protein interactions of SYCE3 with SYCE1, SYCE1-SIX6OS1 and SYCE2-TEX12, were determined by co-expression pull-downs. In this, MBP fusions of the core constructs of SYCE1, SYCE1-SIX6OS1 and SYCE2-TEX12 (Davies et al., 2012; Dunne and Davies, 2019b; Sanchez-Saez et al., 2020) were co-expressed with His- and GST-tagged SYCE3 constructs. For each construct, 3 l cultures were grown. Cells were lysed by sonication in 30 ml 20 mM Tris pH 8.0, 500mM KCl, and lysate clarified by high-speed centrifugation. The supernatant was applied to 6 ml amylose resin (NEB) at 4 °C. After washing with 30 ml 20 mM Tris pH 8.0, 150 mM KCl, 2 mM DTT, bound complexes were eluted in 20 mM Tris pH 8.0, 150 mM KCl, 2 mM DTT, 30 mM D-maltose. Sample concentrations were normalised and analysed by SDS-PAGE.

### Co-purification interaction studies

The relative stability of SYCP1 and SYCE3 protein complexes was assessed by co-expression followed by stringent purification to determine their co-purification or dissociation. MBP-fusions of SYCP1 constructs were co-expressed with His-tagged SYCE3 constructs and were grown in 4 litre cultures, lysed and applied to 8ml amylose resin in 20 mM Tris pH 8.0, 500 mM KCl, 2 mM DTT. Amylose elutions were purified by anion exchange chromatography (HiTrap Q HP, Cytiva) and size exclusion chromatography (HiLoad^TM^ 16/600 Superdex^TM^ 200, Cytiva) in 20 mM Tris pH 8.0, 150 mM KCl, 2 mM DTT. MBP-SYCP1 elution fractions were pooled, and samples of equal concentrations were analysed by SDS-PAGE.

### SYCP1-SYCE3 gel-filtration interaction studies

To analyse the SYCP1αNcore-SYCE3 interaction, 50 µl protein samples were prepared corresponding to SYCP1αNcore, SYCE3, the purified SYCP1αNcore-SYCE3 complex and a 1:1 mixture of SYCP1αNcore and SYCE3, with each component at 235 µM. To analyse the SYCP1αN-SYCE3 interaction, 50 µl protein samples were prepared corresponding to SYCP1αN, SYCE3 and mixtures of SYCP1αN and SYCE3 at 1:0.5, 1:5 and 1:10 molar ratios, in which the concentration of SYCP1αN was 127 µM. Samples were incubated for 1 hour at room temperature and centrifuged at 14000 g at 4°C for 30 minutes. Size exclusion chromatography was performed using a Superdex™ 200 Increase 10/300 GL column in 20 mM Tris pH 8.0, 150 mM KCl, 2 mM DTT at 0.5 ml/min. Elution fractions were analysed by SDS-PAGE.

### SYCP1-SYCE3 gel-filtration assembly assays

The SYCP1αNcore-SYCE3 complex was mixed with SYCE3 at a 10-fold molar excess (92 µM complex with 920 µM SYCE3 and SYCE3 W41E Y44E). SYCP1αN-SYCE3 complex was mixed with a 10-fold molar excess of SYCE3 (93 µM complex with 930 µM SYCE3 and SYCE3 W41E Y44E). Mixed samples and individual components were incubated for 1 hour at 30°C and centrifuged at 14000 g at 4°C for 30 minutes. Size exclusion chromatography was performed using a Superdex™ 200 Increase 10/300 GL (Cytiva) column in 20 mM Tris pH 8.0, 150 mM KCl, 2 mM DTT at 0.5 ml/min. Elution fractions were analysed by SDS-PAGE.

### Circular dichroism (CD)

Far-UV CD spectra were measured using a Jasco J-810 spectropolarimeter (Institute for Cell and Molecular Biosciences, Newcastle University). Wavelength scans were measured at 4°C between 260 and 185 nm at 0.2 nm intervals at using a quartz cuvette, 0.2 mm pathlength (Hellma), with protein samples at 0.2-0.4 mg/ml in 10 mM Na_2_HPO_4_ pH 7.5, 150 mM NaF. For each sample, nine measurements were recorded, averaged and buffer corrected for conversion to mean residue ellipticity ([θ]) (x1000 deg.cm^2^.dmol^-1^.residue^-1^) with deconvolution carried out using the Dichroweb CDSSTR algorithm (http://dichroweb.cryst.bbk.ac.uk). CD thermal melts were recorded between 5°C and 95°C, at intervals of 0.5°C with a 1°C per minute ramping rate, and measured at 222 nm. Protein samples were measured at 0.1 mg/ml in 20 mM Tris pH 8.0, 150 mM KCl, 2 mM DTT, in a 1 mm pathlength quartz cuvette (Hellma) and data plotted as % unfolded after conversion to MRE ([θ]_222,x_-[θ]_222,5_)/([θ]_222,95_-[θ]_222,5_) with melting temperatures determined as the temperature at which the samples are 50% unfolded.

### Isothermal calorimetry (ITC)

ITC data were collected using a Malvern iTC200 instrument at 30°C on samples that had been dialysed overnight into 25 mM Tris pH 8.0, 250 mM KCl, 2 mM DTT, centrifuged at 14000 g at room temperature for 5 minutes, and degassed. SYCE3 (400 µM) was injected into the sample cell containing SYCP1αNcore (70 µM). 19 injections of 4 second duration and 2 µl volume (and an initial injection of 0.2 µl and 0.4 seconds) were performed at intervals of 240 seconds with stirring at 750 rpm. Data were integrated, fitted and plotted using *NITPIC* (Keller et al., 2012), SEDPHAT (Zhao et al., 2015) and GUSSI (https://www.utsouthwestern.edu/labs/mbr/software/), following reported protocols (Brautigam et al., 2016).

### Microscale thermophoresis (MST)

Proteins were labelled in 10 mM HEPES pH 8.0, 150 mM NaCl using the Monolith NT Protein Labelling Kit RED (NanoTemper Technologies) according to the manufacturer’s protocol. Labelled proteins were kept at a constant concentration indicated in the respective figure legends. The unlabelled interacting partner was titrated in 1:1 dilutions. Measurements were performed in premium treated capillaries (NanoTemper Technologies) on a Monolith NT.115 system (NanoTemper Technologies) and excitation and MST power were set at 40 %. Laser on and off times were set at 5 and 30 seconds, respectively.

### Size-exclusion chromatography multi-angle light scattering (SEC-MALS)

The oligomeric state of protein samples was determined by SEC-MALS analysis of protein samples at 5-20 mg/ml in 20 mM Tris pH 8.0, 150 mM KCl, 2 mM DTT. Samples were loaded at 0.5 ml/min onto a Superdex™ 200 Increase 10/300 GL (GE Healthcare) column with an ÄKTA™ Pure (GE Healthcare). The column outflow was fed into a DAWN® HELEOS™ II MALS detector (Wyatt Technology), and then an Optilab® T-rEX™ differential refractometer (Wyatt Technology). ASTRA® 6 software (Wyatt Technology) was used to collect and analyse SEC-MALS data, using Zimm plot extrapolation with a 0.185 ml/g dn/dc value to determine molecular weights from eluted protein peaks.

### Size-exclusion chromatography small-angle X-ray scattering (SEC-SAXS)

SEC-SAXS experiments were performed on beamline B21 at Diamond Light Source synchrotron facility (Oxfordshire, UK). Protein samples at concentrations >5 mg/ml were loaded onto a Superdex™ 200 Increase 10/300 GL size exclusion chromatography column (GE Healthcare) in 20 mM Tris pH 8.0, 150 mM KCl at 0.5 ml/min using an Agilent 1200 HPLC system. The column elution passed through the experimental cell, with SAXS data recorded at 12.4 keV, detector distance 4.014 m, in 3.0 s frames. ScÅtter 3.0 (http://www.bioisis.net) was used to subtract, average the frames and carry out the Guinier analysis for the radius of gyration (*Rg*), and *P(r)* distributions were fitted using *PRIMUS* (P.V.Konarev, 2003). *Ab initio* modelling was performed using DAMMIF (Franke and Svergun, 2009) imposing P1 or P2 symmetry (as indicated) and 30 independent runs were averaged and displayed as DAMFILT envelopes.

### Yeast two-hybrid (Y2H)

Sequences corresponding to human SYCP1 (1-811, 101-783, 1-362, 101-362, 1-206) and SYCE3 (1-88) were cloned into pGBKT7 vectors (Clontech) and human sequences for SYCP1 (1-811), SYCE3 (1-88), SYCE1 (1-351), SYCE2 (1-218), TEX12 (1-123) and SIX6OS1 (1-587) were cloned into pGADT7 vectors (Clontech). The Matchmaker™ Gold system (Clontech) was used for Y2H analysis, using manufacturer’s instructions. Yeast transformations were performed by the standard PEG/ssDNA/LiAc protocol, with the Y187 strain transformed with pGBKT7 vectors and Y2H Gold strain transformed with pGADT7 vectors. The two strains were mated in 0.5 ml 2xYPDA at 30°C, 50 r.p.m, by mixing respective single colonies. Mated cultures were pelleted and resuspended in 0.5xYPDA for plating onto SD/-Trp/-Leu to select for mated colonies and also onto SD/-Trp/-Leu/-Ade/-His with X-α-gal to detect mated colonies that have activated the ADE1, HIS3 and MEL1 reporter genes. Plates were incubated for 5 days at 30°C before imaging.

### Transmission electron microscopy (TEM)

TEM experiments were performed using a Philips CM100 TEM (Electron Microscopy Research services, Newcastle University). SYCE2-TEX12 samples at 3 mg/ml were incubated with a two-fold molar excess of SYCE3 and were applied to carbon-coated grids, washed and then negatively stained with 0.1% v/v uranyl acetate for imaging.

### Protein structure analysis

Molecular structures images were generated using the PyMOL Molecular Graphics System, Version 2.4 Schrödinger, LLC.

### CRISPR/Cas9 Gene Editing

*Syce3* mutant mice were generated by Alt-R CRISPR (Quadros et al., 2017) using a paired nickase design to minimise off-target mutations (Shen et al., 2014). Guide RNA complexes were prepared by annealing 1:1 molar ratios of crRNA (oligos Syce3_20092 or Syce3_20053; Supplementary Table 1) and tracrRNA (IDT). CBAB6F1 female mice (Charles River) were superovulated with 5 IU pregnant mare serum followed by 5 IU human chorionic gonadotophin 42-48 hours later, then mated with CBAB6F1 males. Zygotes were isolated at E0.5 and Alt-R CRISPR reagents (20 ng/µL Alt-R S.p. Cas9 D10A nickase V3 (IDT), 10 ng/µL each guide RNA complex, 20 ng/µL total ssDNA repair oligo in 10 mM Tris pH 7.5, 0.1 mM EDTA) microinjected into the cytoplasm. Zygotes were cultured overnight in KSOM, then transferred to the oviduct of pseudopregnant recipient females. The resulting pups were genotyped from ear clips by sequencing the PCR products obtained using primers Syce3_O2F and Syce3_O2R (Supplementary Table 1). The WY repair oligo introduces W41E and Y41E amino acid mutations into *Syce3* and includes two silent mutations within the nickase PAM sites (Supplementary Table 1). The control PAM repair oligo only contains the two silent PAM site mutations. Adult F0 animals were culled by cervical dislocation and tissues dissected in PBS for analysis. For the silent PAM site mutations, two *Syce3^PAM/PAM^* homozygous animals were obtained directly from the CRISPR/Cas9 injections, additional homozygous animals were then generated by breeding. Matings with male or female mice carrying the *Syce3^WY^* allele were not productive.

### Mouse Phenotyping

Adult mice were culled by cervical dislocation at 2-4 months old, and their testes and epididymides dissected in PBS. Testis weights and cauda epididymis sperm counts were obtained as described previously (Ollinger et al., 2008). For testis histology, testes were fixed in Bouin’s fixative, embedded in wax, sectioned, and stained with haematoxylin and eosin (Ollinger et al., 2008). Although CRISPR/Cas9 founder animals can exhibit mosaicism (Teboul et al., 2017; Wang et al., 2013), the mouse germline typically originates from only three or four epiblast cells (Soriano and Jaenisch, 1986; Ueno et al., 2009), and we did not detect regions of phenotypic mosaicism in the testes of the animals selected for this study.

### Meiotic Chromosome Spreads

Chromosome spreads were prepared from adult *Syce3* testes as described (Costa et al., 2005). Chromosome spreads were stained with antibodies as described (Crichton et al., 2017). Primary antibodies were mouse anti-SYCP3 (Abcam #ab97672, 1:500), rabbit anti-SYCP1 (Abcam #ab15090, 1:200) and rabbit anti-RAD51 (Millipore #PC 130, 1:500). Slides were mounted using antifade mounting medium (Vectashield, H-1000) and high precision coverslips (Marienfeld).

### Widefield Fluorescent Imaging

Widefield epifluorescent images were acquired for a single plane using a Zeiss Axioplan II fluorescence microscope with a Photometrics Coolsnap HQ2 CCD camera, and multiple z-planes using a Zeiss AxioImager M2 fluorescence microscope with a Photometrics Prime BSI CMOS camera. Image capture was performed using Micromanager (Version 1.4), z-stacks were deconvolved in Huygens Essential and maximum intensity projected, and all images were analysed in ImageJ.

### Super-Resolution Imaging

Three dimensional SIM images were captured with a Nikon N-SIM microscope with an Andor iXon 897 EMCCD camera (Andor technologies, Belfast UK). Consistent capture parameters were used for given antibody combinations. Chromosome spreads that extended beyond the field of view were captured as multiple images with 15% overlap (Supplementary Figure 13), then stitched together using Nikon NIS-Elements. Maximum intensity projections were taken forward for further analysis. Custom pipelines in ImageJ, Python and R were used to quantitatively analyse the SIM images.

### Quantitative Image Analysis

Binary masks were generated in ImageJ by manual thresholding of antibody-stained channels, and individual nuclear territories by drawing a region of interest around DAPI staining. Masks were converted into labelmaps for focal staining patterns. Downstream analysis was performed in Python3 and R.

Focal labels were shuffled within the nuclear territory by randomly assigning new centroid coordinates to each focus within the nuclear space, ensuring that the edges of each focus territory did not overlap one another or exceed the nuclear boundary. For calculation of mean fluorescence intensity within foci, the mean nuclear background signal from the area not assigned to foci was first subtracted to control for background variation.

### Animal Ethics

Experiments involving animals were conducted in line with institutional and national ethical and welfare guidelines and regulations. The experiments described in this study were approved by the University of Edinburgh animal welfare and ethics review board and performed under authority of UK Home Office licences PP3007F29 and PB0DC8431.

### Accession codes and data availability

All data are available from the corresponding authors upon reasonable request.

## Acknowledgements

We thank Diamond Light Source and the staff of beamline B21 (proposals sm14435, sm15580, sm15897 and sm15836). We thank H. Waller for assistance with CD data collection, and V. A. Jatikusumo and M. Ratcliff for work in the early stages of this project. We thank A. Wheeler, M. Pearson and L. Murphy in the MRC HGU advanced imaging resource for help and guidance with imaging and image analysis, the Edinburgh Super-Resolution Imaging Consortium for super-resolution imaging, and the University of Edinburgh Bioresearch & Veterinary Services for mouse husbandry. We thank Wendy Bickmore, Javier Caceres, Cova Vara and Adele Marston for critically reviewing the manuscript. I.R.A. and J.H.C. are supported by MRC University Unit grant MC_UU_00007/6. O.R.D. is a Wellcome Senior Research Fellow (Grant Number 219413/Z/19/Z).

## Author contributions

J.M.D., O.M.D. and L.J.S. performed biochemical and biophysical experiments. J.H.C. performed mouse phenotyping, developed imaging analysis pipelines and analysed imaging. P.D. performed CRISPR/Cas9 injections. J.L. performed mouse genotyping. O.R.D. and I.R.A. analysed data, designed experiments and wrote the manuscript.

---

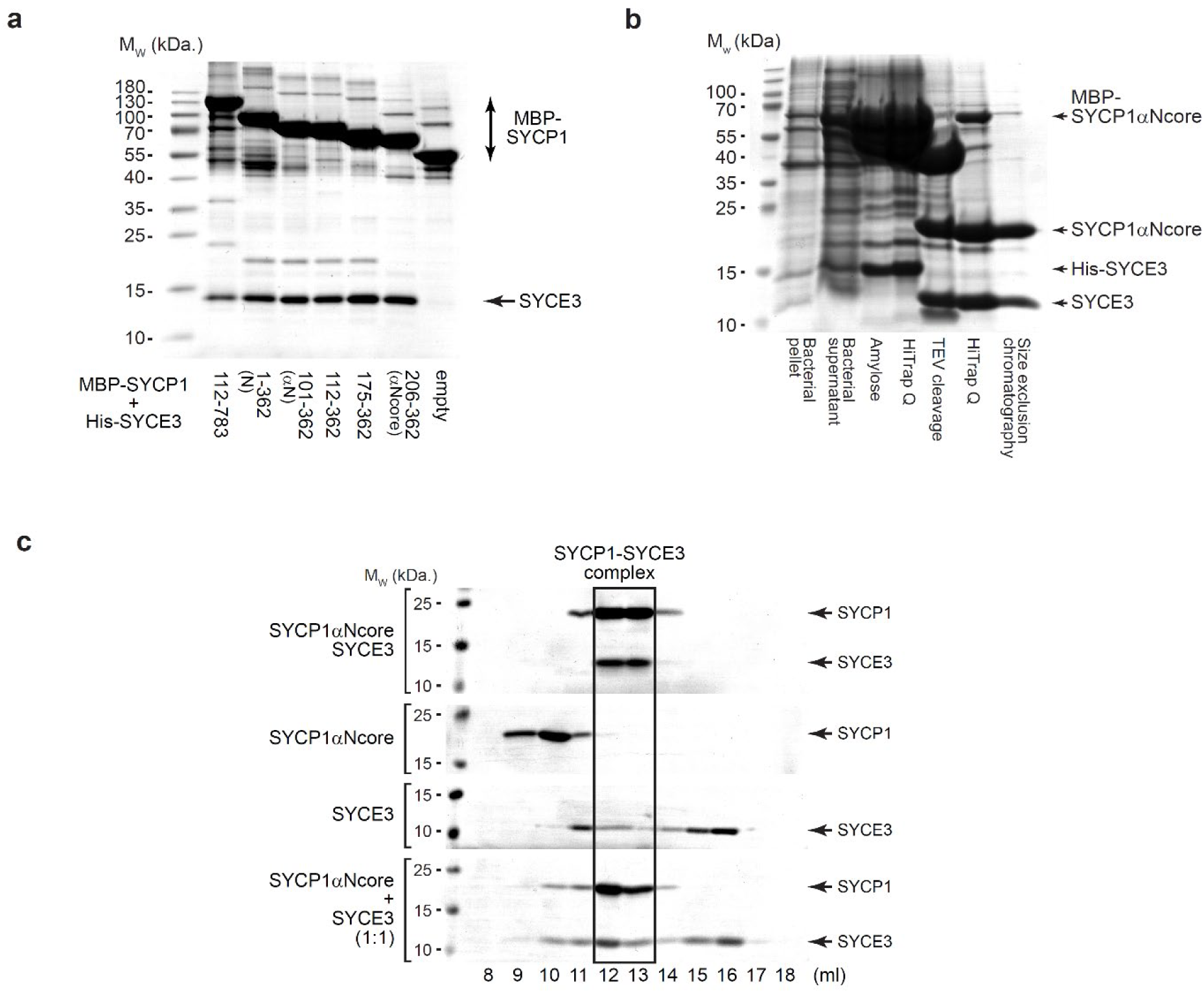
SYCP1 forms a high-affinity complex with SYCE3. (**a**) Uncropped gel of Figure 1e, corresponding to amylose pull-down of His-SYCE3 following recombinant co-expression with MBP-SYCP1 constructs (as indicated) and free MBP (empty). (**b**) Recombinant co-expression and co-purification of SYCP1αNcore-SYCE3 through amylose and anion exchange chromatography, followed by TEV cleavage to remove N-terminal expression tags, with subsequent anion exchange and size exclusion chromatography. (**c**) SDS-PAGE of elution fractions corresponding to the size-exclusion chromatography analysis shown in Figure 1f.

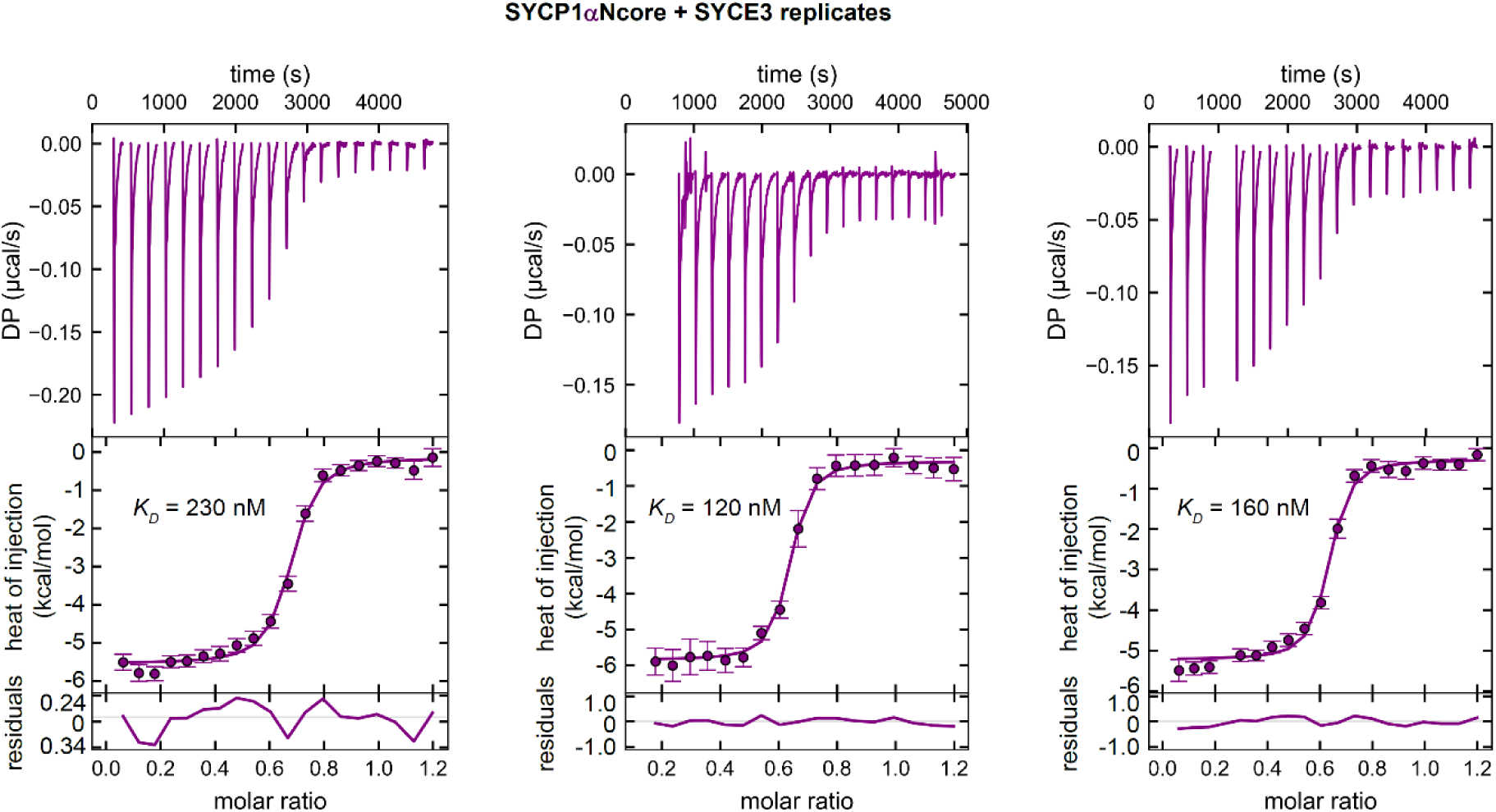
ITC and analysis of SYCP1-SYCE3 complex formation. Replicates of ITC analysis of SYCE3 titrated into SYCP1αNcore, corresponding to Figure 1h, showing injections (top), fit (middle) and residuals (bottom), with individual replicate apparent affinities of 230 nM, 120 nM and 160 nM.

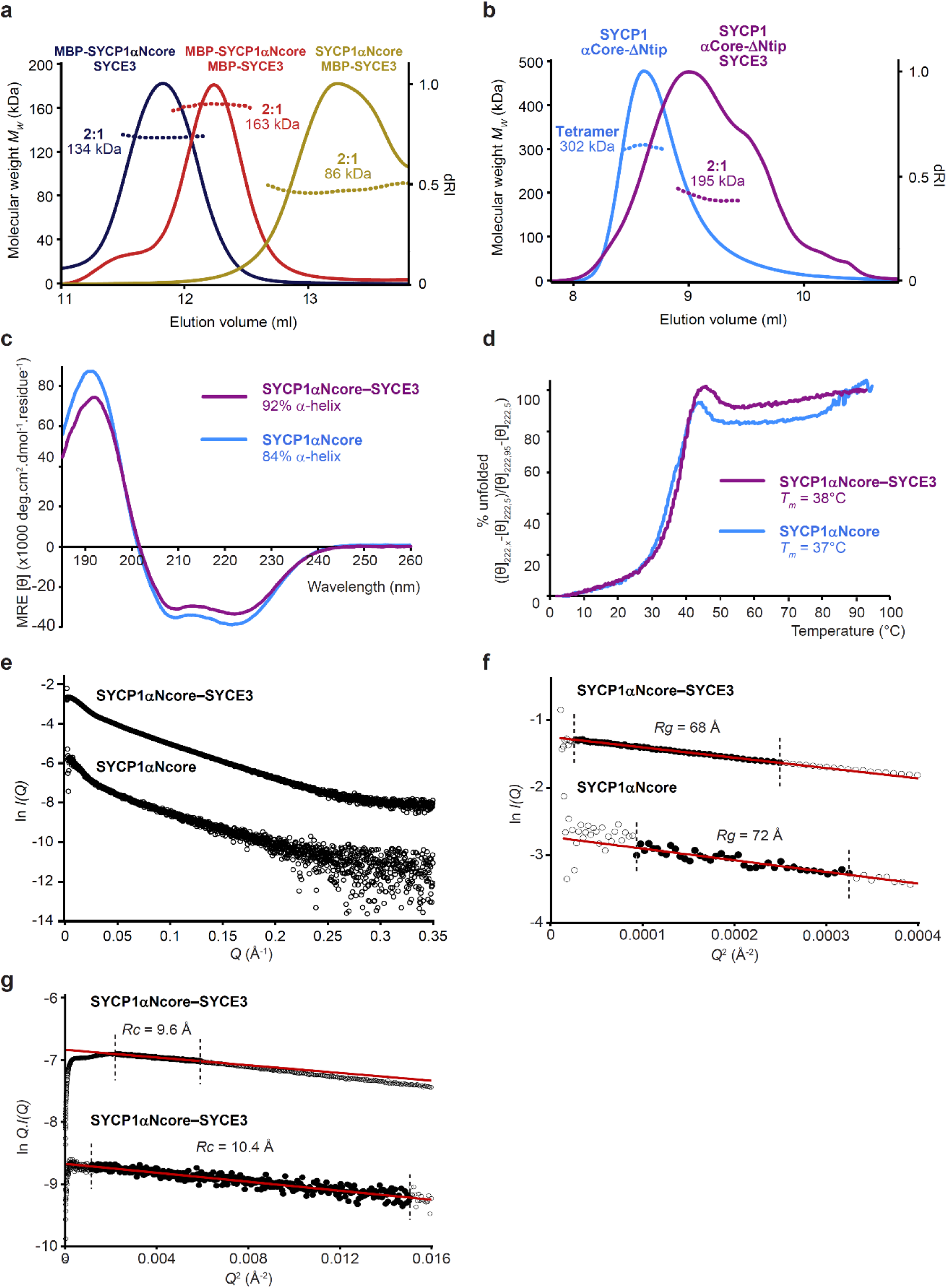
Structure of the SYCP1αNcore-SYCE3 complex. (**a**) SEC-MALS analysis of MBP-SYCP1αNcore-SYCE3 (blue), MBP-SYCP1αNcore-MBP-SYCE3 (red), SYCP1αNcore-MBP-SYCE3 (yellow), revealing 2:1 complexes of 134 kDa, 163 kDa and 86 kDa, respectively (theoretical – 134 kDa, 175 kDa and 94 kDa). (**b**) SEC-MALS analysis of SYCP1 αCore-ΔNtip in isolation and in complex with SYCE3, demonstrating a 302 kDa tetramer and 195 kDa 2:1 complex, respectively (theoretical – 320 kDa and 171 kDa). (**c**) Far UV CD spectra and (**d**) CD thermal denaturation of SYCP1αNcore-SYCE3 (purple) and SYCP1αNcore (blue). (**c**) Secondary structure composition was estimated through deconvolution of spectra with data fitted at normalised rms deviation values of 0.006 and 0.001, respectively. (**d**) Thermal denaturation recorded for SYCP1αNcore-SYCE3 and SYCP1αNcore as % unfolded based on the helical signal at 222 nm; melting temperatures were estimated at 38°C and 37°C, respectively. (**e-g**) SEC-SAXS analysis. (**e**) Scattering intensity plots, (**f**) Guinier analysis to determine the radius of gyration (Rg) with linear fits shown in black (Q.Rg values were < 1.3) and (**g**) Guinier analysis to determine the radius of gyration of the cross-section (Rc) (Q.Rc values were < 1.3) for SYCP1αNcore-SYCE3 and SYCP1αNcore. Corresponding *P(r)* distributions and *ab initio* models are shown in Figure 2b.

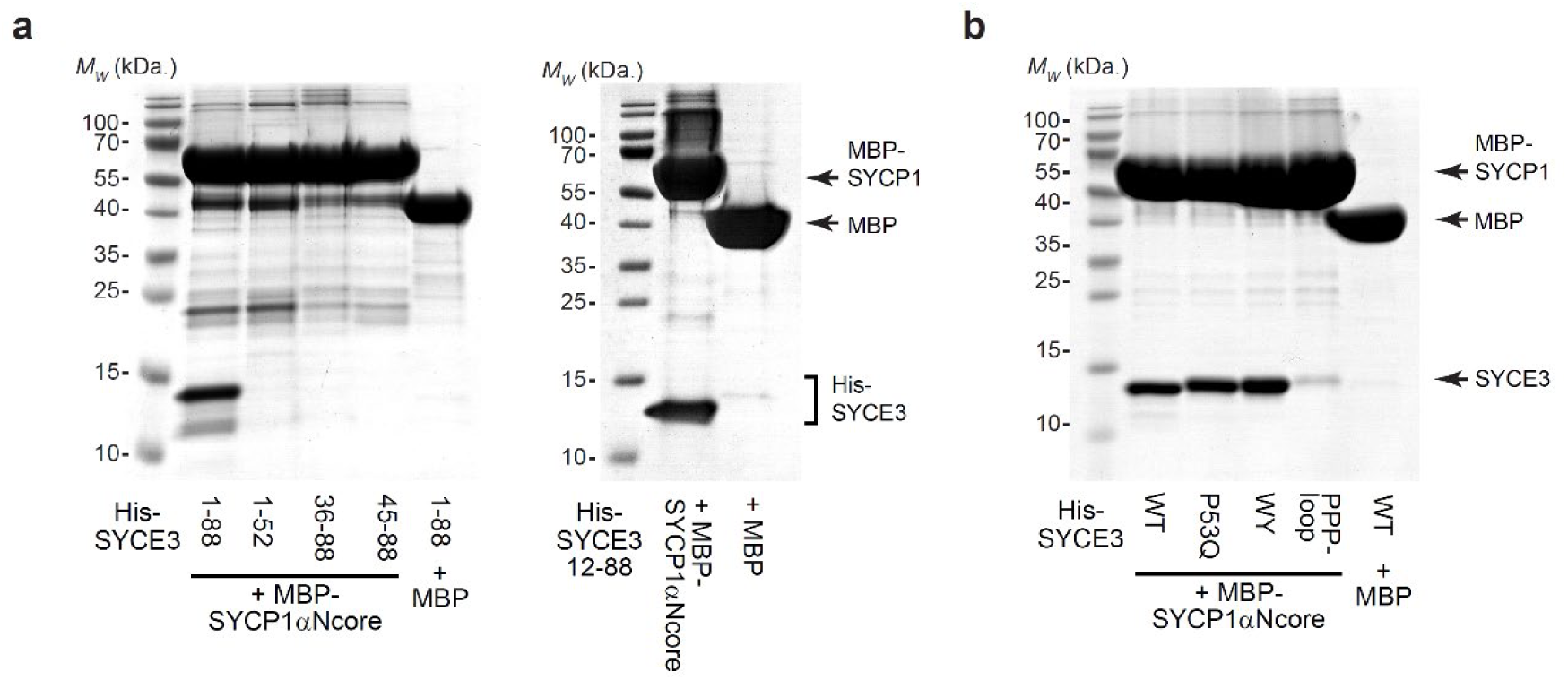
Pull-downs of SYCP1-SYCE3 mutant complexes. (**a,b**) SYCE3-binding analysis through co-expression with MBP-SYCP1 or free MBP and co-purification by amylose, ion exchange and size-exclusion chromatography using (**a**) SYCP1αNcore and SYCE3 wild-type (WT), P53Q, W41E Y44E (WY) and PPP-loop, and (**b**) SYCP1αNcore and SYCE3 truncations.

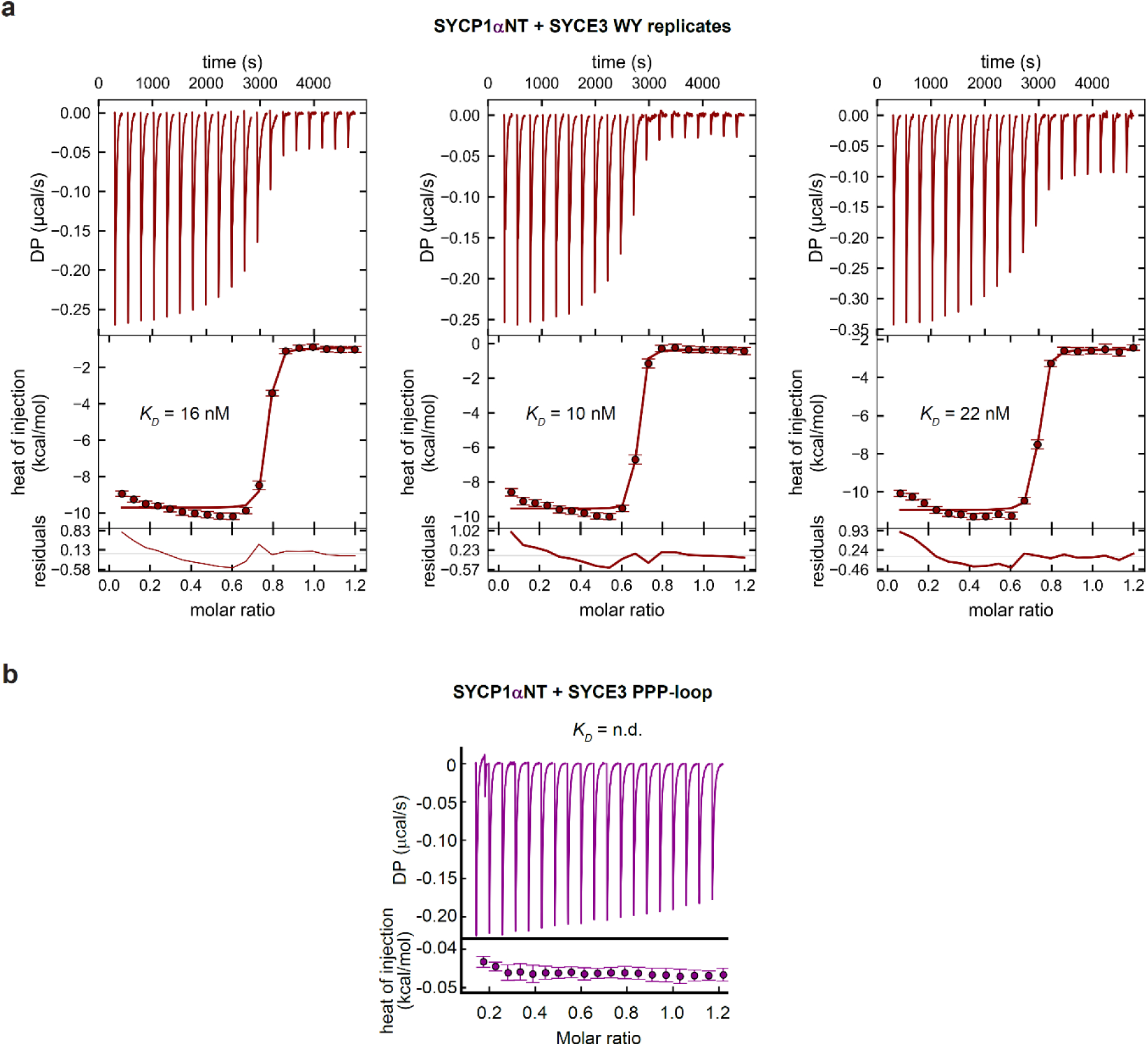
ITC and analysis of SYCP1-binding by SYCE3 WY and PPP-loop. (**a**) Replicates of ITC analysis of SYCE3 WY titrated into SYCP1αNcore, corresponding to Figure 2g, showing injections (top), fit (middle) and residuals (bottom), with individual replicate apparent affinities of 16 nM, 10 nM and 22 nM. (**b**) ITC of SYCE3 PPP-loop titrated into SYCP1αNT, in which no interaction was not observed and the binding affinity was not determined (n.d.).

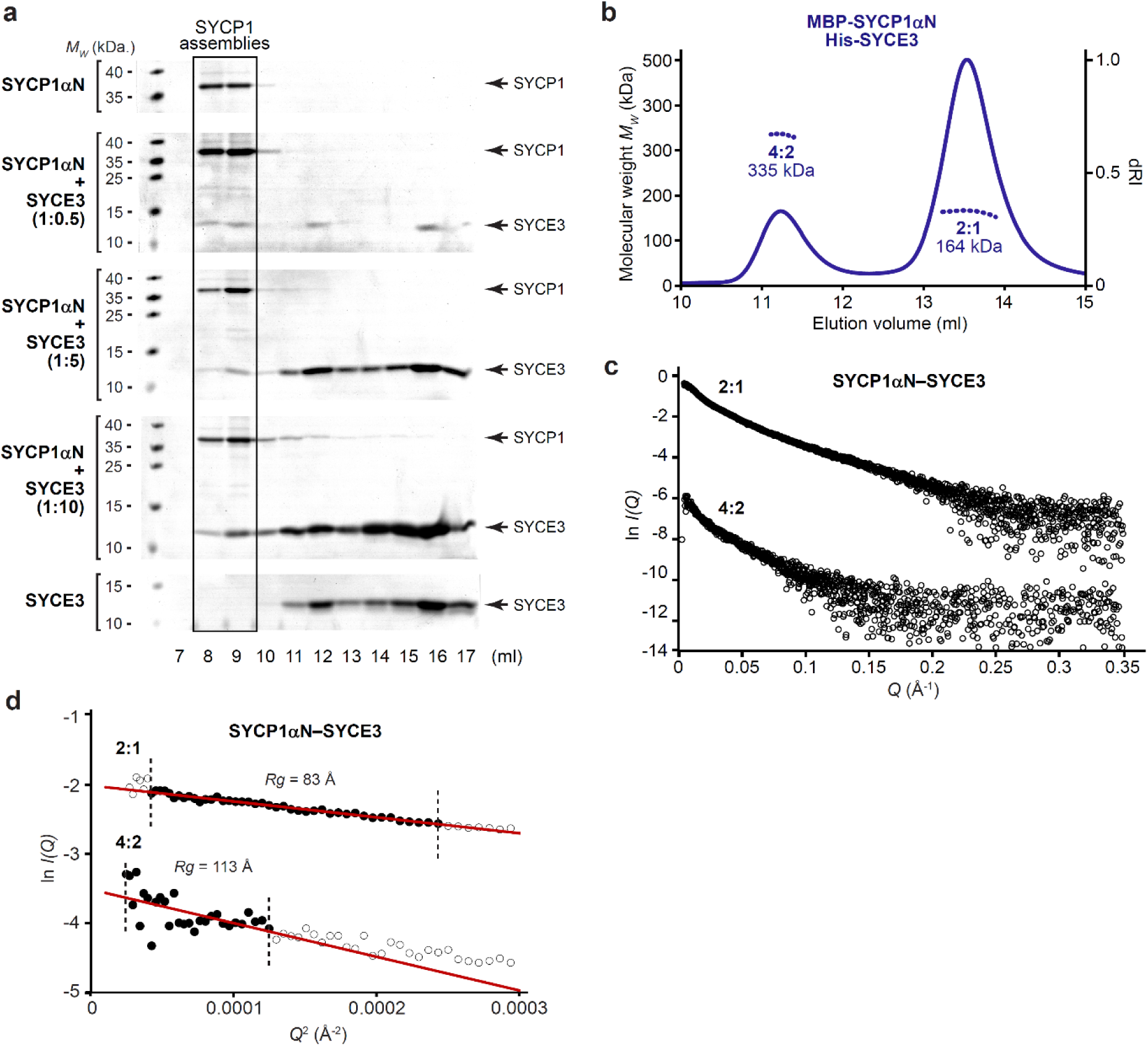
Structure of the SYCP1αN-SYCE3 complex. (**a**) SDS-PAGE of size-exclusion chromatography elution fractions of SYCP1αN upon incubation with SYCE3 at stoichiometric ratios of 1:0.5, 1:5 and 1:10; free SYCP1αN and SYCE3 are shown for comparison. (**b**) SEC-MALS analysis (using a Superose 6 increase 10/300 GL column) of MBP-SYCP1αN-His-SYCE3 revealing 2:1 and 4:2 species of 164 kDa and 335 kDa, respectively (theoretical – 160 kDa and 319 kDa). (**c,d**) SEC-SAXS analysis. (**c**) Scattering intensity plots and (**d**) Guinier analysis to determine the radius of gyration (Rg) with linear fits shown in black (Q.Rg values were < 1.3) for SYCP1αN-SYCE3 2:1 and 4:2 complexes. Corresponding *P(r)* distributions and *ab initio* models are shown in Figure 3c,d.

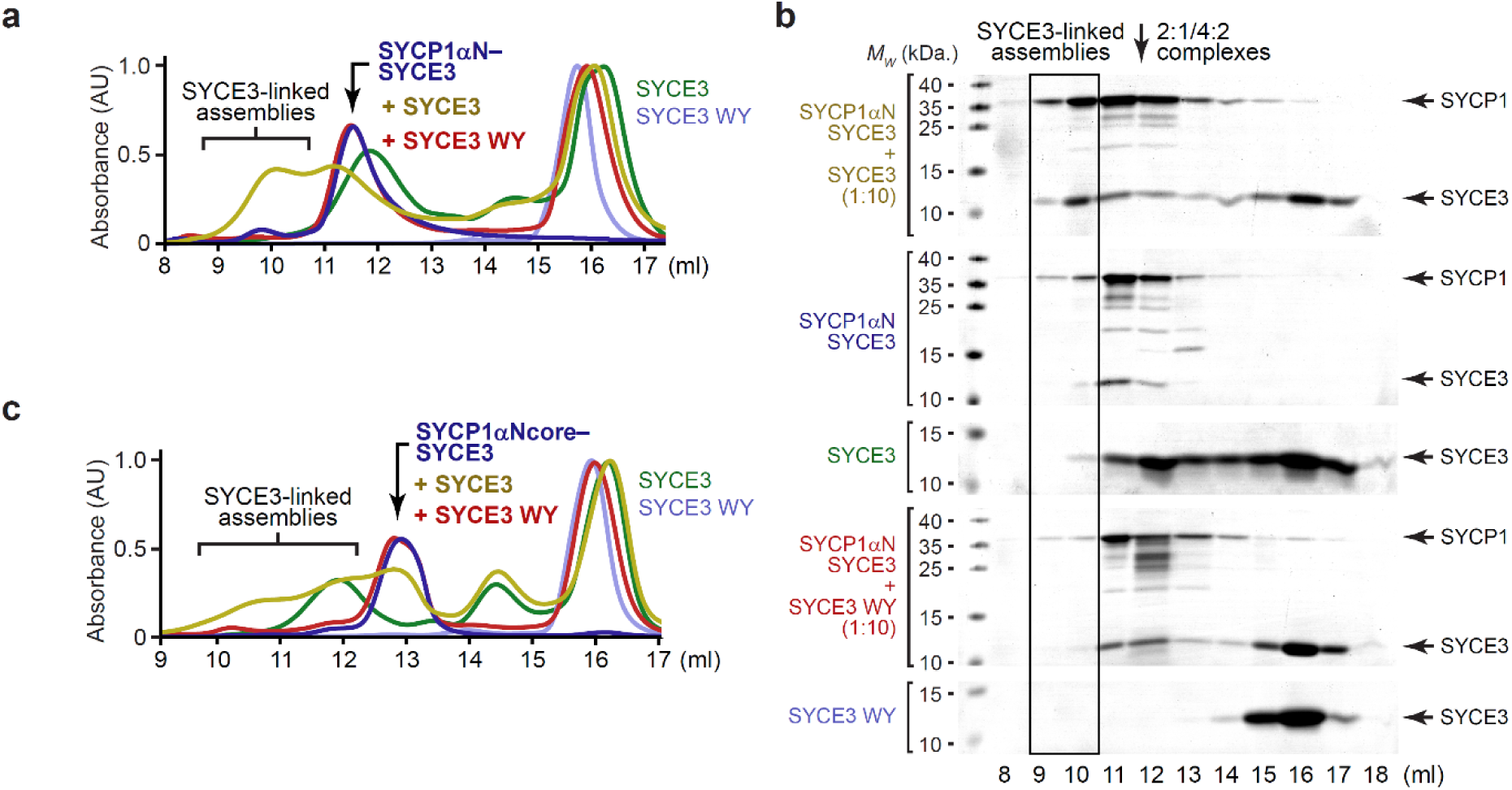
SYCP1-SYCE3 integrated lattice formation through SYCE3 self-assembly. (**a**) Size-exclusion chromatography of (**a,b**) SYCP1αN-SYCE3 and (**c**) SYCP1αNcore-SYCE3 upon incubation with a 10-fold stoichiometric excess of SYCE3 wild-type or WY, corresponding to Figure 4f-h. (**a,c**) UV absorbance (280 nm) chromatograms normalised to the same maximum peak height shown in Figure 3f,g with additional chromatograms for free SYCE3 wild-type and WY.

**Supplementary Figure 8.**
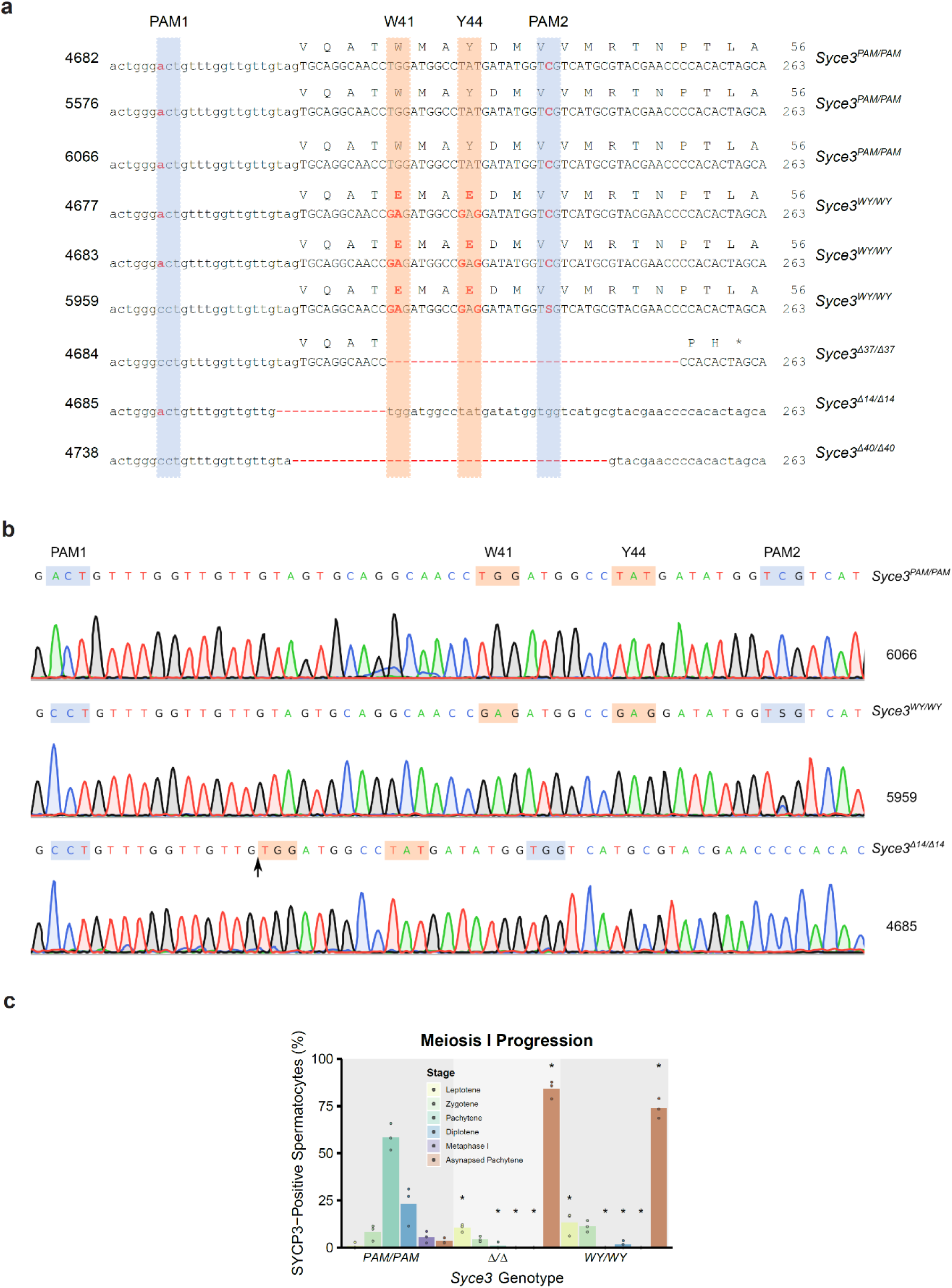
*Syce3* mutant allele sequences and meiotic phenotypes. (**a**) *Syce3* nucleotide and predicted protein sequences of mice used in this study. Mouse IDs are shown on the left and the genotype group on the right. Animal 5959 in the *Syce3^WY/WY^* group did not incorporate the silent C:A mutation in PAM1 and was mosaic/heterozygous for the silent G:C mutation in PAM2. For the *Syce3^Δ/Δ^* group the number of nucleotides deleted is indicated in the allele name. The PAM sequences are highlighted with blue boxes, the W41 and Y44 sequences with orange boxes. (**b**) Chromatograms showing examples of sequencing the *Syce3* locus from mice with the indicated genotypes and IDs. Mice that lacked potential mosaicism/heterogeneity at the W41/Y44 codons were used in this study. (**c**) Percentage of SYCP3-positive spermatocytes at the indicated stage of meiosis I in *Syce3* chromosome spreads based on SYCP3 and SYCP1 immunostaining (Figure 4f). Mean percentages for each genotype are indicated by the bars, percentages from individual animals are indicated by filled circles. Asterisks indicate a significant difference (p<0.05, Student’s t-test, n=3) relative to *Syce3^PAM/PAM^* controls. Summary statistics are in Supplementary Table 2.

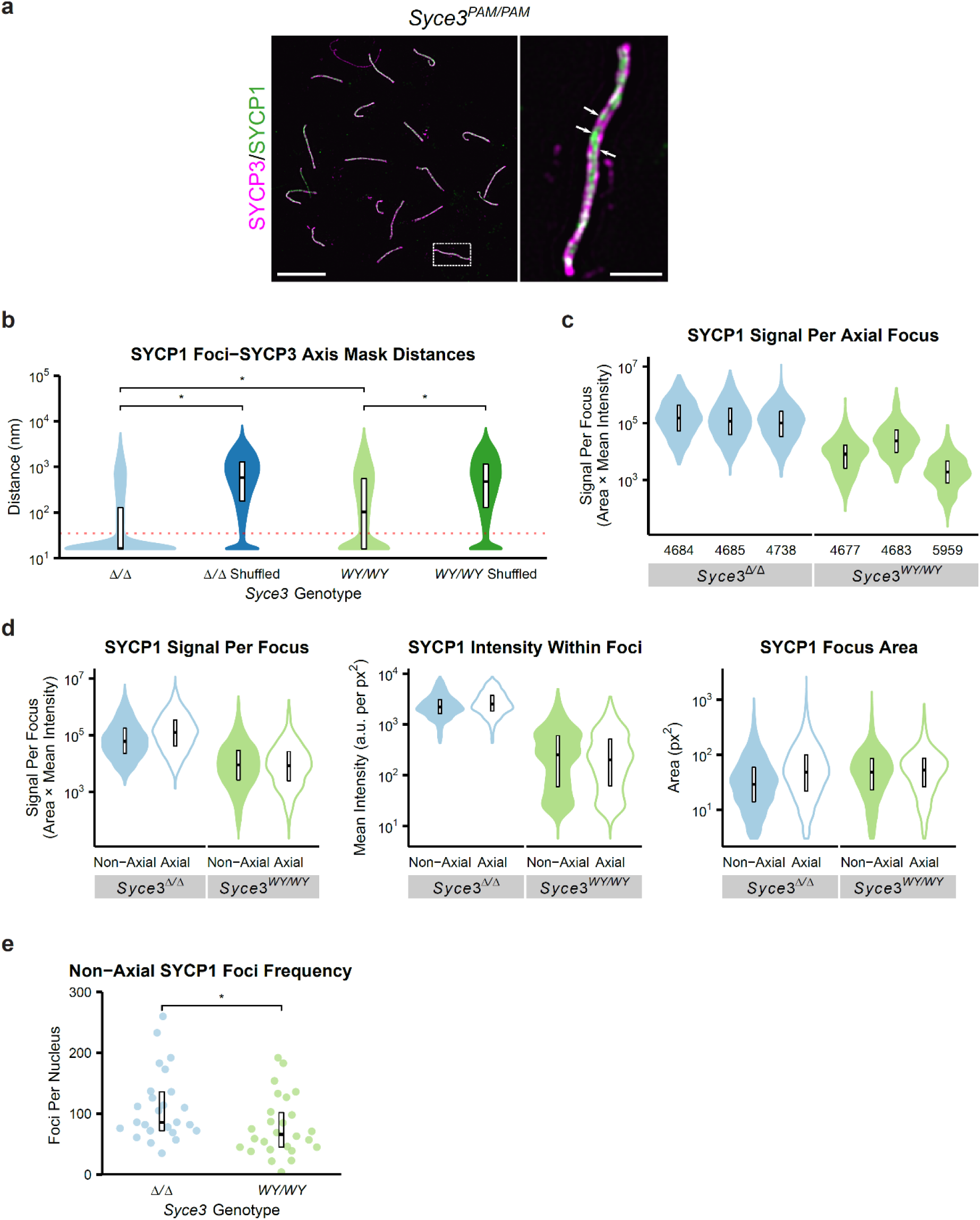
Quantitative analysis of SYCP1 foci in *Syce3* spermatocytes. (**a**) SIM images of pachytene *Syce3^PAM/PAM^* meiotic chromosome spreads immunostained for SYCP3 (magenta) and SYCP1 (green). Scale bars, 10 µm for low magnification images, 1 µm for enlarged regions. Patches where SYCP1 extends linearly along one axis are indicated with arrows. The spread shown is the same as in Figure 5a (**b**) SYCP1 foci-SYCE3 axis mask distances. Distances from the centroid of each SYCP1 focus to the nearest point on the SYCP3 axis mask are shown alongside distances from shuffled datasets obtained by assigning all SYCP1 foci in each nucleus to a random nuclear location for twenty iterations. The red dotted horizontal line represents the 35 nm threshold distinguishing axial and non-axial foci. Crossbars represent quartiles; *, p< 0.01 (Mann-Whitney U test, medians are 16, 585, 104 and 473 nm); 3 animals analysed for each *Syce3* genotype. Summary statistics are in Supplementary Table 2. (**c**) Total SYCP1 signal in each axial SYCP1 focus as shown in Figure 5d, with data segmented for individual animals. Crossbars represent quartiles; medians are 152975, 116615, 101019, 8067, 24188 and 1855 arbitrary units); mouse IDs are shown below each dataset. Summary statistics are in Supplementary Table 2. (**d**) Violin plots showing the anti-SYCP1 immunostaining signal per focus, intensity within foci and focus area for axial and non-axial SYCP1 foci in *Syce3^Δ/Δ^* and *Syce3^WY/WY^* spermatocytes. Crossbars represent quartiles. Median SYCP1 signals per focus: 60101, 123441, 9061 and 8485 arbitrary units. Median SYCP1 intensities within foci; 2205, 2525, 255, and 203 arbitrary units per px^2^. Median areas; 29, 48, 48 and 53 nm^2^. 3 animals analysed for each *Syce3* genotype. Summary statistics are in Supplementary Table 2. (**e**) Non-axial SYCP1 foci frequencies in asynapsed pachytene *Syce3^Δ/Δ^* and *Syce3^WY/WY^* spermatocytes. *, p< 0.05 (Mann-Whitney U test, medians are 86 and 66 foci); 3 animals analysed for each *Syce3* genotype. Summary statistics are in Supplementary Table 2.

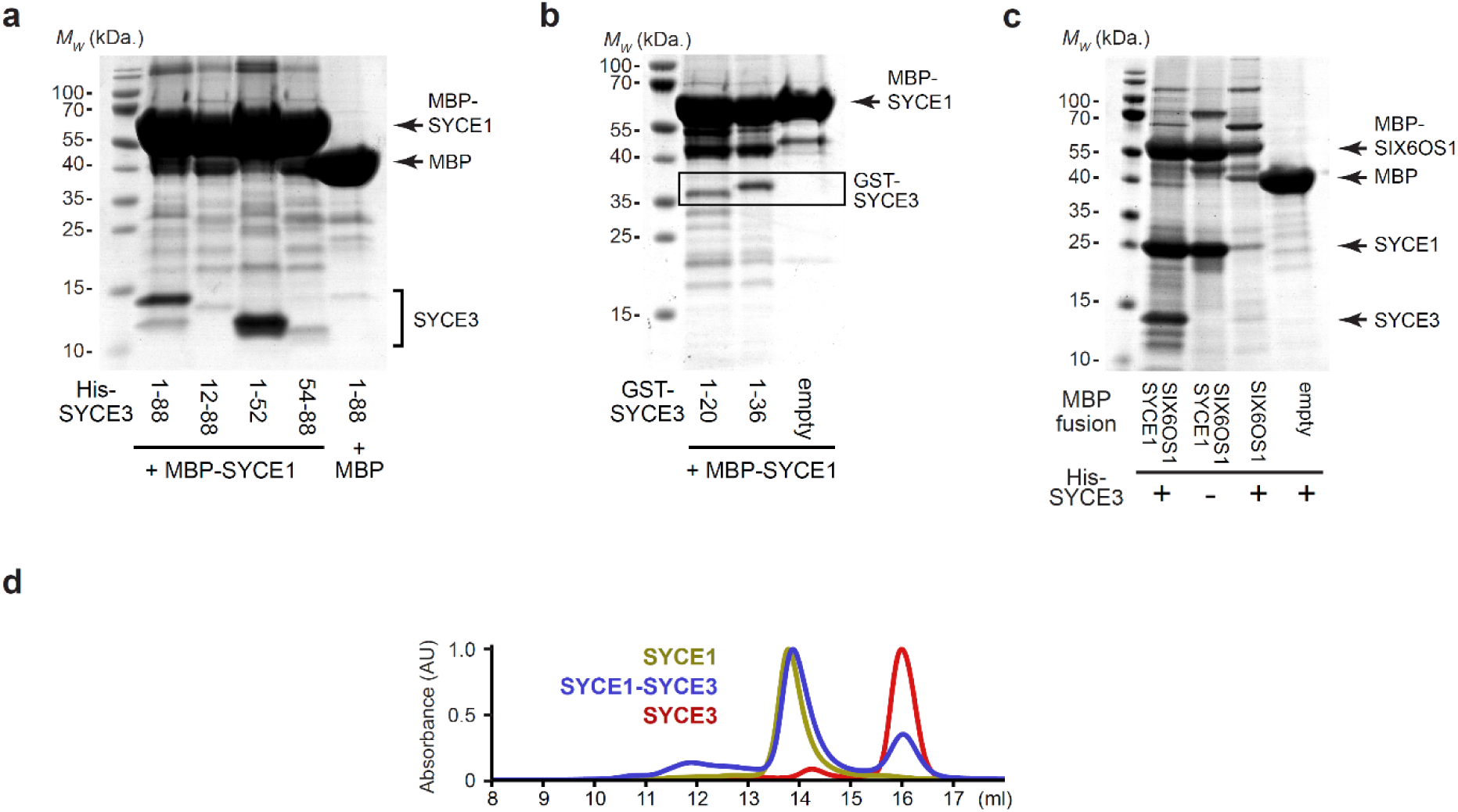
SYCE3 interacts with the SYCE1-SIX6OS1 complex. (**a**) Amylose pull-downs of His-SYCE3 and following recombinant co-expression with MBP-fusions of SYCE1core, SYCE1core-SIX6OS1N, SIX6OS1N, and free MBP, corresponding to Figure 6a-c. (**b**) Size-exclusion chromatography of co-expressed and co-purified SYCE1core-SYCE3, with UV chromatograms of SYCE1core-SYCE3 (blue), SYCE1core (yellow) and SYCE3 (red), corresponding to Figure 6e.

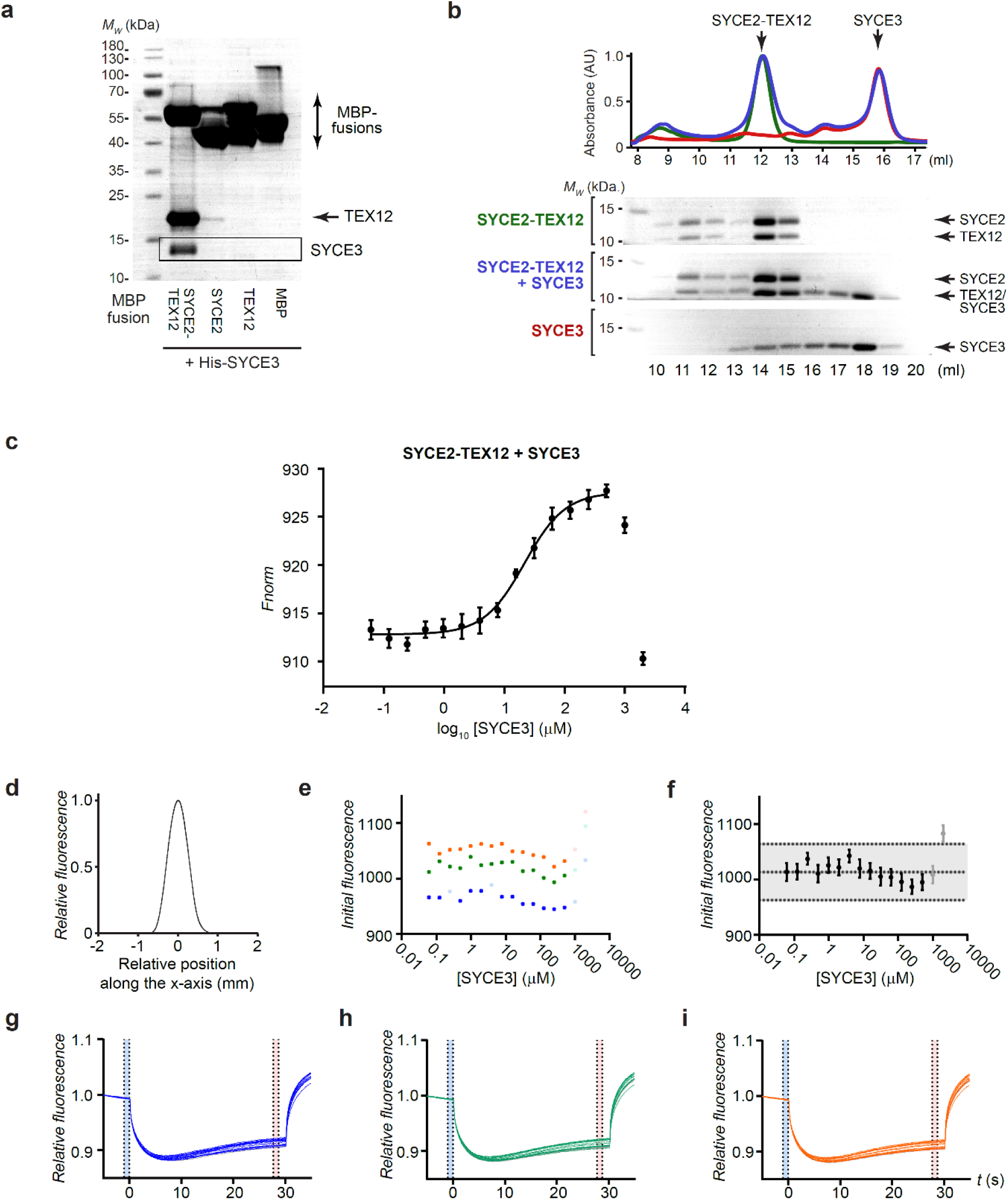
SYCE3 interacts with the SYCE2-TEX12 complex. (**a**) Uncropped gel of Figure 6d, corresponding to amylose pull-downs of His-SYCE3 following co-expression with MBP-fusions of the structure cores of SYCE2-TEX12, SYCE2 and TEX12, and free MBP. (**b-h**) MST analysis corresponding to Figure 6f. (**b**) Full dataset in which the final two datapoints were excluded from analysis. (**c**) Overlaid capillary scans. (**d,e**) Initial fluorescence for three data series (blue, yellow, green) displayed as (**d**) individual data series and (**e**) mean ± SEM (n=3). (**f-h**) Relative fluorescence for each of the three data series. (**i**) Size-exclusion chromatography of SYCE2-TEX12 core (green), SYCE3 (red) and an equimolar mixture of SYCE2-TEX12 core and SYCE3 (blue), shown as UV absorbance (280 nm) chromatograms normalised to the same maximum peak height and SDS-PAGE of elution fractions.

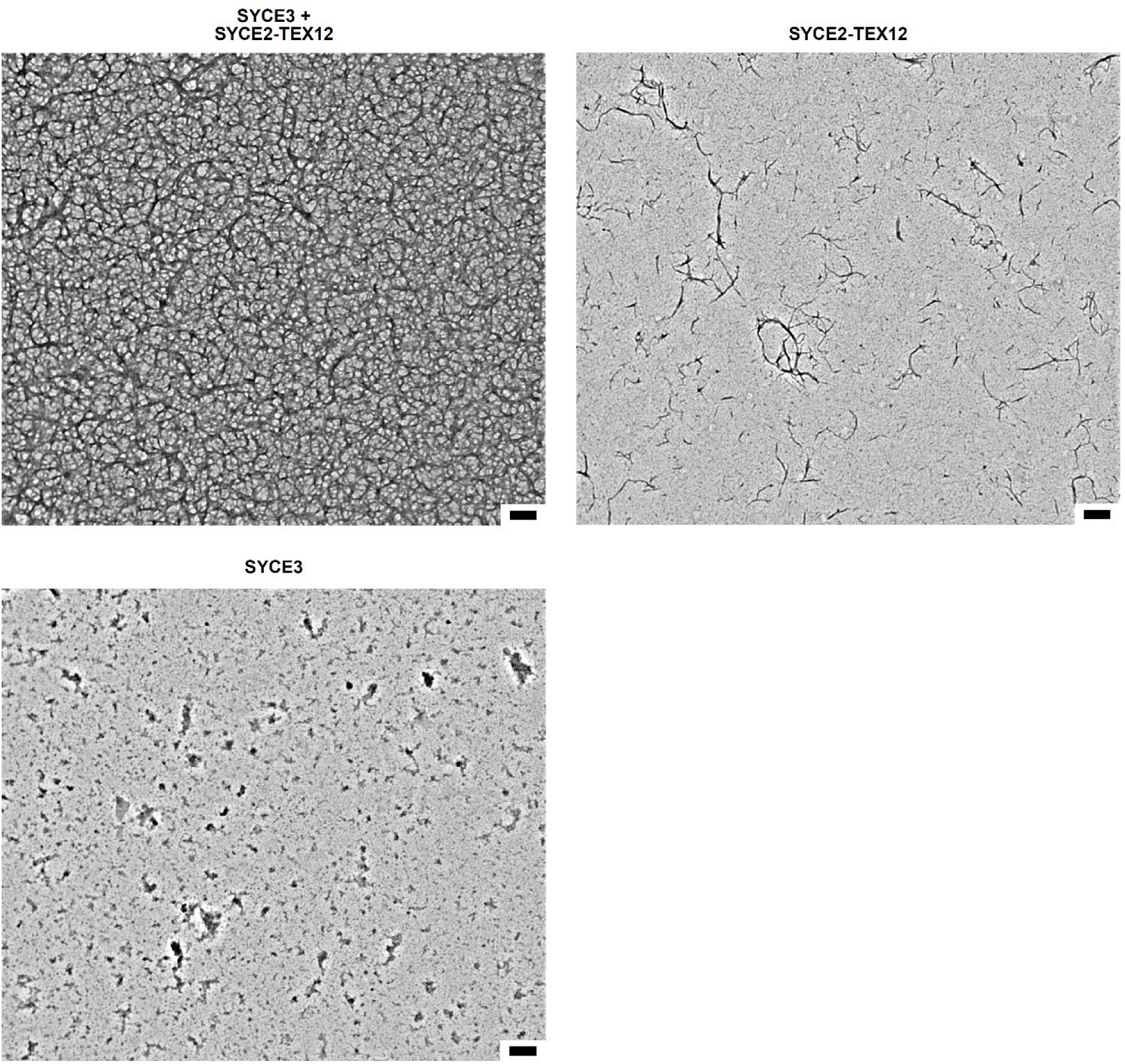
Electron microscopy analysis of SYCE2-TEX12 following incubation with a two-fold excess of SYCE3. Full panels corresponding to Figure 6g. Scale bar, 200 nm.

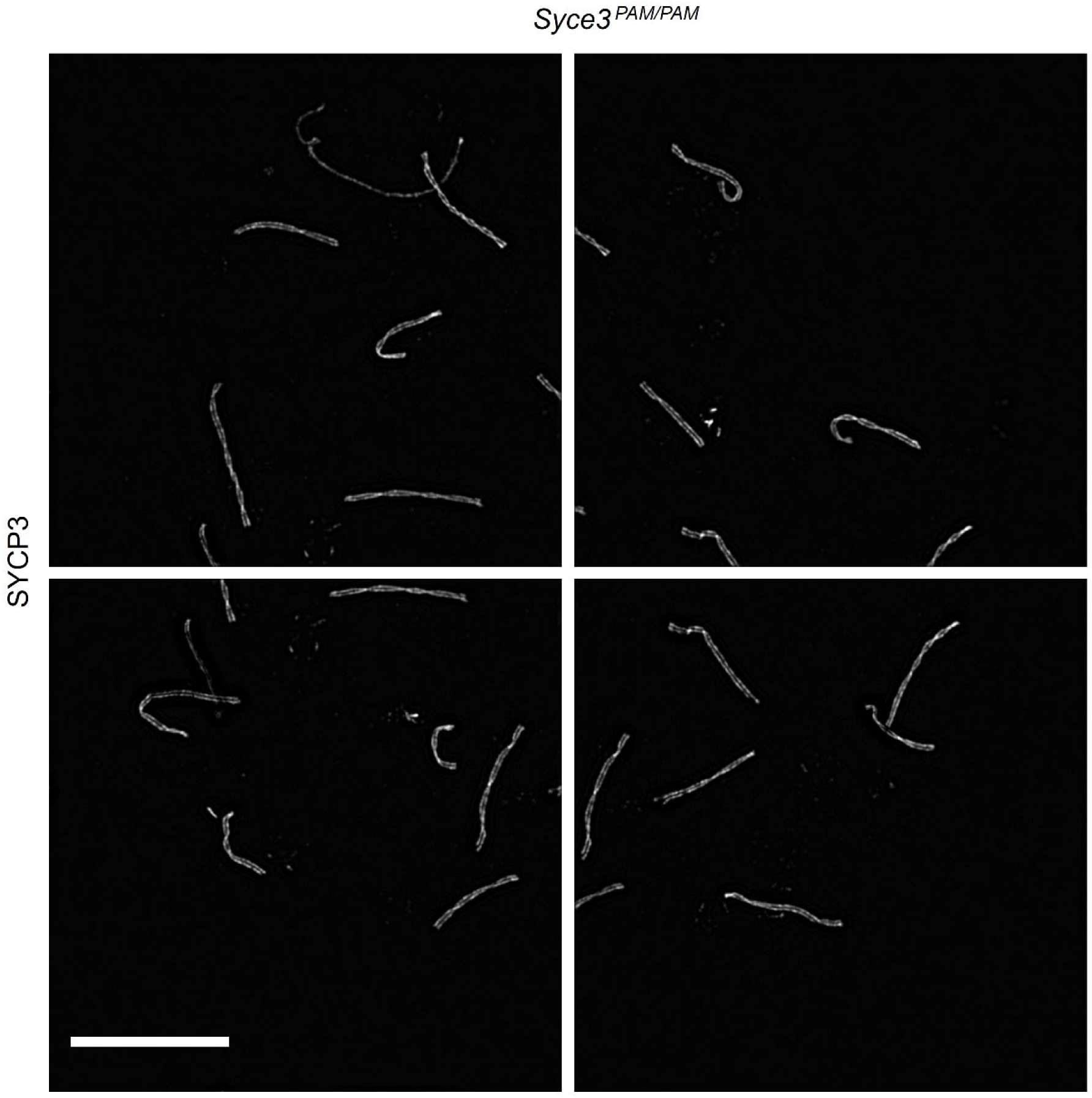
Overlapping images used to capture chromosome spreads. *Syce3^PAM/PAM^* images used to capture the chromosome spread shown in Figure 5b and Supplementary Figure 9a. Scale bar 10 µm. Overlapping images were taken with 15% overlap and stitched using algorithms in Nikon NIS-Elements.

**Supplementary Table 1.**
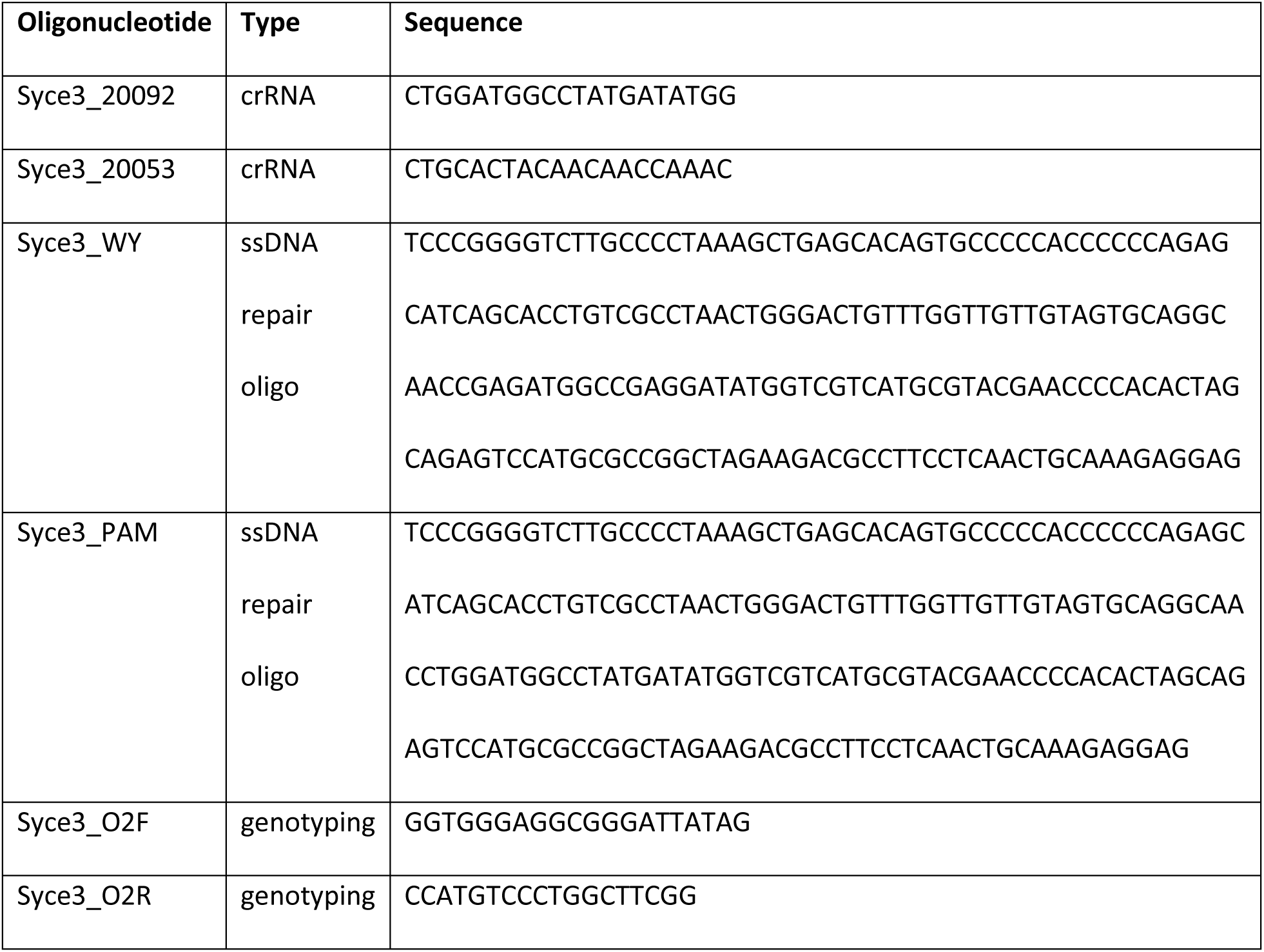
Oligonucleotides used in this study.

